# HRS1/HHOs GARP transcription factors and reactive oxygen species are regulators of Arabidopsis nitrogen starvation response

**DOI:** 10.1101/164277

**Authors:** Alaeddine Safi, Anna Medici, Wojciech Szponarski, Amy Marshall-Colon, Sandrine Ruffel, Frédéic Gaymard, Gloria Coruzzi, Benoît Lacombe, Gabriel Krouk

**Affiliations:** Laboratoire de Biochimie et Physiologie Moléculaire des Plantes, UMR5004 CNRS/INRA/SupAgro/UM, Institut de Biologie Intégrative des Plantes ‘Claude Grignon’, Place Pierre Viala, 34060 Montpellier, France; New York University, Department of Biology, Center for Genomics & Systems Biology, 12 Waverly Place, New York, N.Y. 10003, USA

## Abstract

Plants need to cope with strong variations in the nitrogen content of the soil solution. Although many molecular actors are being discovered concerning how plants perceive NO_3_^-^ provision, it is less clear how plants recognize a lack of Nitrogen. Indeed, following N removal plants activate their Nitrogen Starvation Response (NSR) being characterized in particular by the activation of very high affinity nitrate transport systems (NRT2.4, NRT2.5) and other sentinel genes such as GDH3. Here we show using a combination of functional genomics (via *TF perturbation*) and molecular physiology studies, that the GARP Transcription Factors (TFs) belonging the HHO sub-family are important regulators of the NSR through two potential mechanisms. First, HHOs directly repress *NRT2.4* and *NRT2.5* high-affinity nitrate transporters. Genotypes affected in HHO genes (mutants and overexpressors) display modified high-affinity nitrate transport activities opening interesting perspectives in biotechnology applications. Second, we show that Reactive Oxygen Species (ROS) are important to control NSR in wild type plants and that HRS1 and HHO1 overexpressors are affected in their ROS content, defining a potential feedforward branch of the signaling pathway. Taken together our results define two new classes of molecular actors in the control of NSR including ROS and the first transcription factors to date. This work (i) opens perspectives on a poorly understood nutrient related signaling pathway, and (ii) defines targets for molecular breeding of plants with enhanced NO_3_^-^ uptake.

## Introduction

The fertilization of crops with nitrogen (N) is a key requirement for global food production systems, sustaining the world’s population and ensuring food security. As N is *the* key rate-limiting nutrient in plant growth, understanding the factors that limit N-use efficiency (NUE) will have particular relevance (Han et al., 2015). Inefficient NUE by agricultural systems is also responsible for nitrate run-off into water soil and atmosphere. Increased leaching of N into drainage water and the release of atmospheric nitrous oxide and reactive N greenhouse gases, pollutes the troposphere, contributes to global warming, and accelerates the eutrophication of rivers and acidifies soils (Sutton et al., 2011). Thus, understanding the regulation of N transport by plants is likely to contribute to tackle these problems.

As sessile organisms, plants need to adapt to fluctuating nitrogen (N) conditions (Crawford and Glass, 1998; O'Brien et al., 2016). N related adaptations include modification of germination (Alboresi et al., 2005), root and shoot development (Forde and Walch-Liu, 2009; Gruber et al., 2013; Krouk et al., 2010a; O'Brien et al., 2016; Rahayu et al., 2005), flowering time (Castro Marin et al., 2010), transcriptome and metabolome (Krouk et al., 2010a; O'Brien et al., 2016; Scheible et al., 1997; Stitt, 1999; Wang et al., 2004).

Interestingly, one can distinguish between different N-related signaling pathways being activated in response to different N-variation scenarios and being reported by different sets of sentinel genes.

These signaling pathways include the Primary Nitrate Response (PNR) (Hu et al., 2009; Medici and Krouk, 2014) that can be seen when NO_3_^-^-depleted or N-depleted plants are treated with NO_3_^-^. PNR sentinel genes are very quickly (within minutes) (Krouk et al., 2010b) regulated by NO_3_^-^ and include nitrate reductase gene 1 (*NIA1*), nitrite reductase gene 1 (*NIR1*), glucose 6 phosphate dehydrogenase (*G6PDH*), and several others such as the GARP transcription factors *HRS1* (hypersensitive to low Pi-elicited primary root shortening 1 (Liu et al., 2009)) and *HHO1* (HRS1 homologue 1 (Liu et al., 2009)) (Canales et al., 2014; Medici and Krouk, 2014). PNR is probably the most studied and understood N-related signaling pathway for which several molecular actors have been identified so far. These include the NO_3_^-^ transceptor CHL1/NRT1.1/NPF6.3 (Ho et al., 2009), kinases and phosphatase (CIPK8, CIPK23, ABI2, CPK10,30,32) (Ho et al., 2009; Hu et al., 2009; Leran et al., 2015; Liu et al., 2017), and several transcription factors (NLP6/7, TGA1, NRG2, BT2, TCP20, SPL9) (Alvarez et al., 2014; Araus et al., 2016; Castaings et al., 2009; Guan et al., 2017; Krouk et al., 2010b; Marchive et al., 2013; Wang et al., 2009; Xu et al., 2016). Recently a NO_3_^-^-triggered Ca_2_^+^ signal have been shown to be a crucial relay between the NRT1.1 transceptor and the nuclear events controlling PNR (Krouk, 2017; Liu et al., 2017; Riveras et al., 2015).

Long-distance N-related root-shoot-root signals have also been shown to adapt plant development and metabolism to the whole nitrogen status of the plant (Gansel et al., 2001; Li et al., 2014; Ruffel et al., 2011; Ruffel et al., 2015). These long-distance signals can be divided into N-demand signals and N-supply signals, which can be genetically uncoupled (Li et al., 2014; Ruffel et al., 2011; Ruffel et al., 2015). Cytokinin and CEP peptides biosynthesis have been shown to be important to generate the N-demand root-to-shoot-to-root relay necessary to regulate genes and modify root development in conditions where N-supply is heterogeneous (Ohkubo et al., 2017; Ruffel et al., 2011; Ruffel et al., 2015; Tabata et al., 2014).

Finally, another signaling pathway includes N-Starvation Response (NSR). It can be related to the molecular events triggered by a prolonged N-deprivation (Kiba and Krapp, 2016; Krapp et al., 2011; Menz et al., 2016). Sentinel genes of NSR include high affinity (Km~10 μM) NO_3_^-^ transporters NRT2.4 and NRT2.5 (activated to retrieve traces of NO_3_^-^ in the soil), as well as the glutamate dehydrogenase 3 gene (GDH3; hypothesized to be activated to recycle N) (Kiba et al., 2012; Kotur and Glass, 2015; Lezhneva et al., 2014; Marchi et al., 2013). To date, the calcineurin B-like7 displayed an effect on *NRT2.4* and *NRT2.5* gene expression in the context of NSR (Ma et al., 2015). A role of miR169 was shown in the control of NFYA transcription factors in response to NSR with a substantial impact on *NRT2.1* and *NRT1.1* transcriptional regulation (Zhao et al., 2011). However, no proof of the actual role of the NFYA genes themselves on NSR was provided in this work (Zhao et al., 2011). LBD37,38,39 transcription factors over-expression have been shown to impact anthocyanin production on plants with N-deprived status (Rubin et al., 2009) with a potential regulation on nitrate transporters including *NRT2.5* (Rubin et al., 2009). However, no direct regulatory target of LBDs were provided in this previous study (Rubin et al., 2009).

These different N-related signaling pathways are likely to be tightly intertwined but no molecular actors are known to be to play such a role. Here, we show that one of the most strongly and rapidly NO_3_^-^ regulated transcription factor (HRS1) (Canales et al., 2014; Krouk et al., 2010b) and its close homologous GARP TFs (HHOs) (Safi et al., 2017), are involved in NSR regulation. This has very important consequences on high-affinity NO_3_^-^ transport activity, as well on plant growth. We also demonstrate that NSR is abolished by ROS scavenging molecules in wild-type plants (WT) and interferes in HHO-manipulated genotypes, thus defining a two-branched signaling pathway.

## Results

During our recent investigations (Medici et al., 2015) on HRS1 direct targets and their role in the control of root development in response to combination of N and P signals, we remarked a that *NRT2.4* transcript accumulation was repressed upon controlled nuclear entrance (Bargmann et al., 2013; Medici et al., 2015) of the GR:HRS1 fusion protein (see Sup Fig. 1a; (Medici et al., 2015)). We also noticed that *NRT2.4* was over-expressed in the double *hrs1;hho1* mutant (data shown in Sup Fig. 1b). These preliminary results suggested that HRS1 could be a direct regulator of the *NRT2.4* gene.

**Figure 1.**
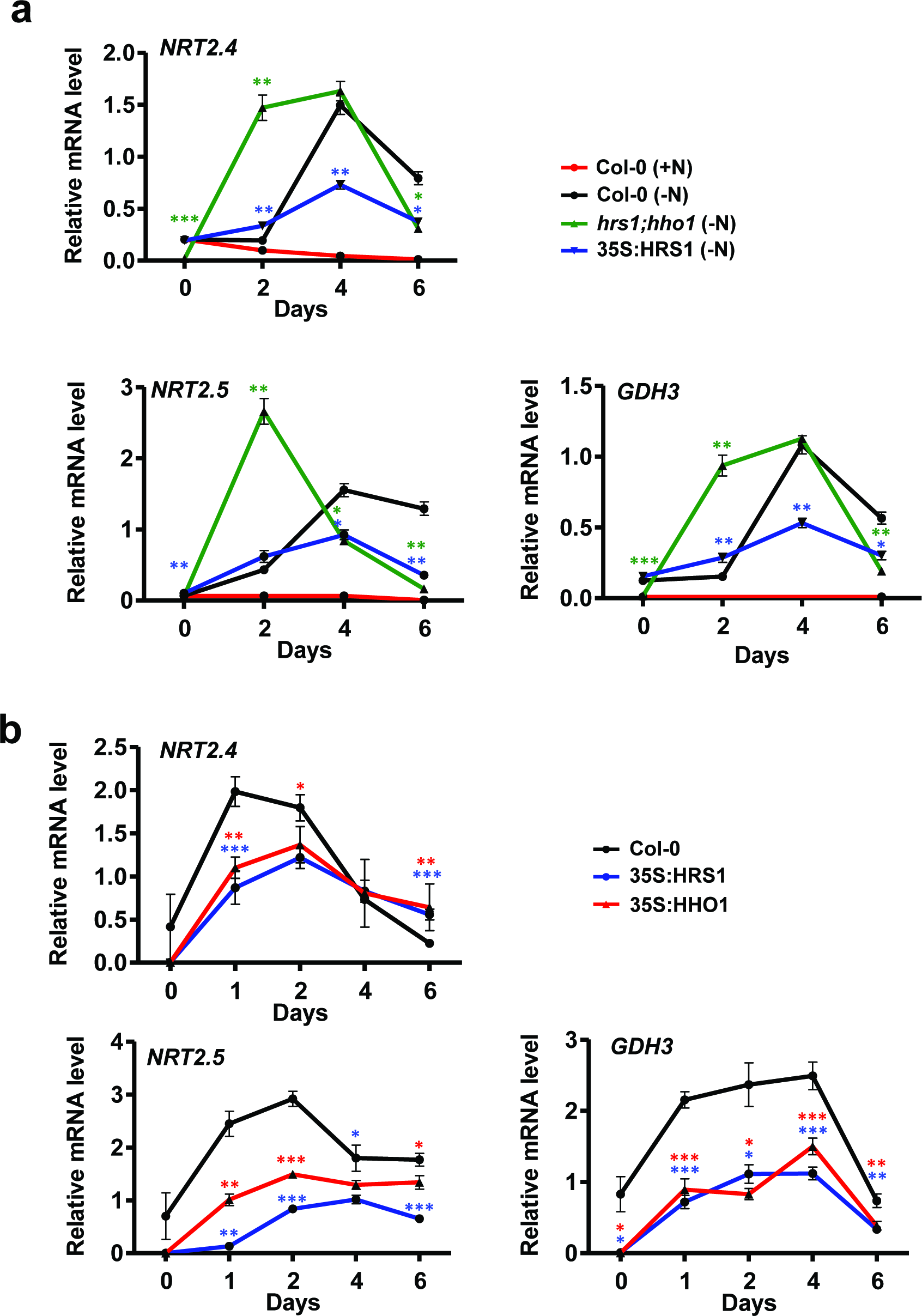
HRS1 and HHO1 are repressors of NSR sentinel genes. **(a)** Root response of *NRT2.4, NRT2.5* and *GDH3* to NSR in Columbia, *hrs1;hho1*, 35S:HRS1 genotypes. Plants are grown in sterile hydroponic conditions on N containing media for 14 days. At time 0, the media is shifted to ‐N conditions for 0, 2, 4, 6 days or +N as a control. **(b)** Root response of *NRT2.4, NRT2.5* and *GDH3* to NSR in Columbia WT, 35S:HRS1, 35S:HHO1 genotypes. Plants are grown in sterile hydroponic conditions on N containing media for 14 days. Then the media is shifted to ‐N conditions for 0, 1, 2, 4, 6 days or +N as a control (the media background is kept unchanged). All transcript levels were quantified by qPCR and normalized to two housekeeping genes (ACT and CLA), values are means ± s.e.m (n= 4). Asterisks indicate significant differences from WT plants (*P<0.05; **P<0.01; ***P<0.001; Student’s t-test).

Moreover, as *NRT2.4* was shown to be a very good marker of the N-starvation response (Kiba et al., 2012), and no regulator was shown to participate in this signaling pathway at the outset of this study, we decided to investigate the role of HRS1 and its close homologous in the NSR response. Since the experiments in Medici et al. (2015) (Sup Fig. 1b) were performed on plants grown in very particular conditions (minus-Phosphate, plus-N), we investigated the behavior of the *hrs1;hho1* mutants and HRS1, HHO1 over-expressors (Sup Fig. 2a) in the specific context of the NSR (transfer of plants from N containing media to N-free media). To consolidate our investigations, we also studied two other genes that are sentinels of the NSR, namely *NRT2.5* (another very high affinity NO_3_^-^ transporter) (Lezhneva et al., 2014), and *GDH3* (Marchi et al., 2013).

**Figure 2.**
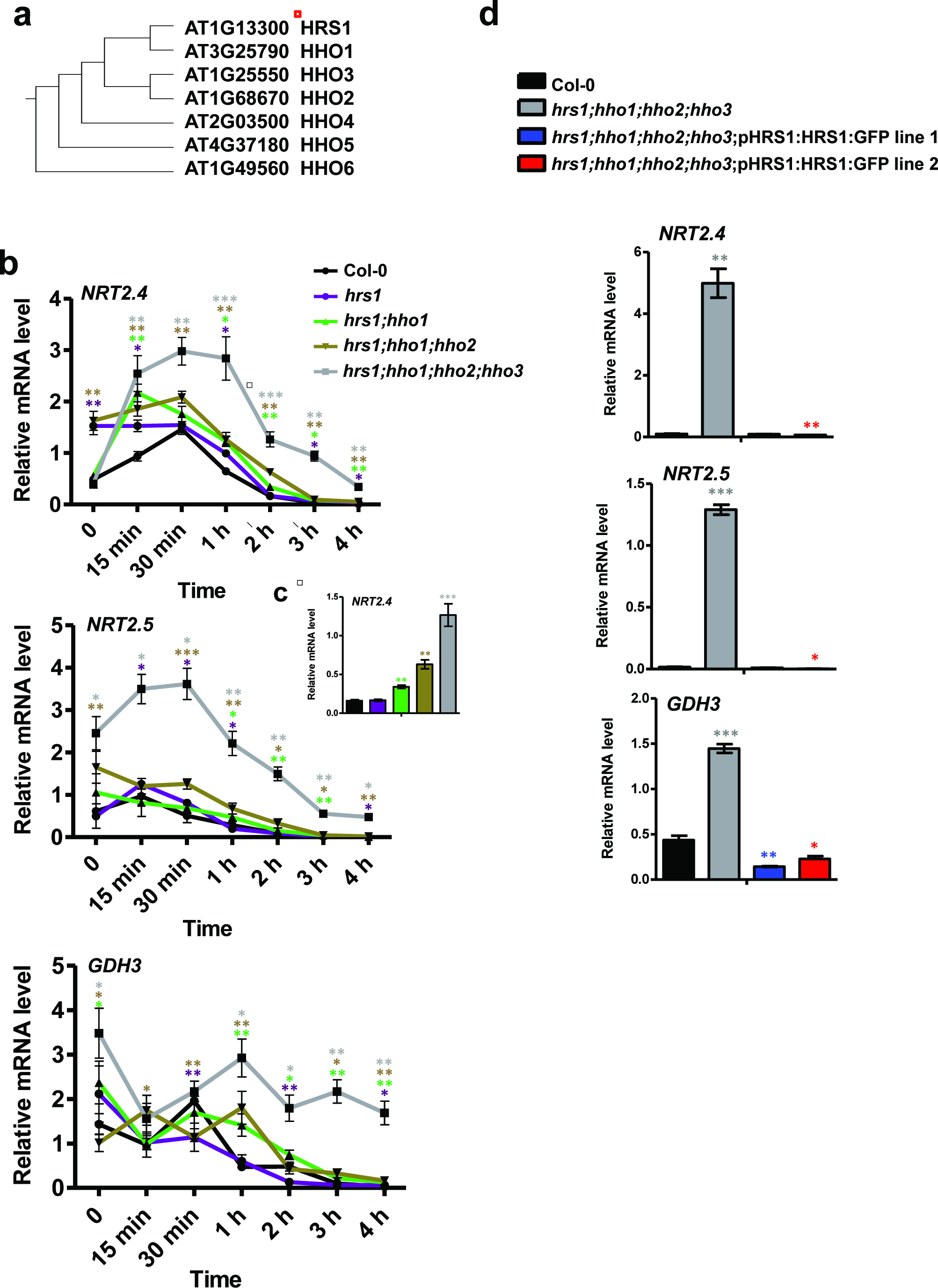
HHO subfamily is involved in repressing NSR sentinels after NO_3_^-^ provision. **(a)** Phylogenetic tree representing the GARP HHO subfamily. The tree was built as described in Safi et al., (2017) using the mafft algorithm (http://mafft.cbrc.jp/alignment/server/, parameters: G-INS-1, BLOSUM62) and drawn with FigTree. **(b)** Root response of *NRT2.4, NRT2.5* and *GDH3* to PNR following N starvation in Columbia WT, *hrs1, hrs1;hho1, hrs1;hho1;hho2, hrs1;hho1;hho2;hho3* genotypes. Plants are grown in sterile hydroponic conditions on full media for 14 days, subjected to N starvation for 3 days and then resupplied with 0.5 mM NH4NO_3_ for 0 (harvest before treatment), 15 min, 30 min, 1h, 2h, 3h, 4h (the media background is kept unchanged). **(c)** pHRS1:HRS1:GFP is sufficient to complement the *hrs1;hho1;hho2;hho3* quadruple mutant. Col, *hrs1;hho1;hho2;hho3*, hrs1;hho1;hho2;hho3;pHRS1:HRS1:GFP line 1 and line 2 (2 independent transformation events) are grown on petri dishes on 0.5 mM of NH_4_NO_3_ for 12 days. Roots are harvested and transcripts are measured by qPCR. All transcript levels were quantified by qPCR and normalized to two housekeeping genes (ACT and CLA), values are means ± s.e.m (n= 4). Asterisks indicate significant differences from WT plants (*P<0.05; **P<0.01; ***P<0.001; Student’s t-test).

### HHOs repress NSR sentinel genes

In the growth context of NSR treatments, as expected (Kiba et al., 2012), we showed that in WT plants NSR is manifested by the strong activation of *NRT2.4*, *NRT2.5*, and *GDH3* genes within the first days of starvation (Fig. 1a). These studies revealed that NSR was diminished in 35S:HRS1 plants. In this experiment, these genes are also affected in the *hrs1;hho1* double mutants. Indeed, in the *hrs1;hho1* genotype, NSR sentinel genes peak earlier and are also repressed at earlier time-points compared to WT (Fig. 1a). Since HRS1 and HHO1 are homologous genes, and because the double mutant has a NSR phenotype, we set up an experiment to compare HRS1 and HHO1 over-expressers side-by-side. This result (Fig. 1b) showed that 35S:HRS1 and 35S:HHO1 display similar molecular phenotypes with a reduction of the NSR response. Taken together, these results suggest that HRS1 and HHO1 are repressors of NSR, likely through the direct regulation of *NRT2.4* and *NRT2.5* loci.

One interesting aspect of the *hrs1;hho1* double mutant, which we observed across the different experiments, was a very erratic response to NSR with the most representative response shown in Figure 1a. In order to explain this phenomenon and to stabilize the phenotype we recalled that HRS1, HHO1 and its paralogs HHO2 and HHO3 (Fig. 2a) are all transcriptionally regulated by PNR (upon plant transfer from a N-depleted media to a NO_3_^-^ containing media) (Krouk et al., 2010b; Wang et al., 2004). To understand the interactions of the HHO paralogs in regulating the NSR, we generated plants with an increasing number of deletions in this gene sub-family (Fig. 2a). We generated the triple *hrs1;hho1;hho2* and quadruple *hrs1;hho1;hho2;hho3* mutants by crossing (characterization in Sup Fig. 2b) and compared them with WT, single and double mutants. The rationale was that; if these HRS/HHO transcription factors (TFs) are indeed repressors of the NSR, they could strongly be regulated by nitrate (in the context of the PNR) in order to quickly stop the NSR response when NO_3_^-^ or Nitrogen is available. We thus tested this hypothesis by subjecting the plants (WT, single, double, triple and quadruple HRS/HHO mutants) to a nitrogen starvation treatment first, then treated them with NO_3_^-^, and then followed the speed of NSR sentinel gene repression (Fig. 2b). In this context, we showed that in WT, consistent to what was observed by Okamoto et al. (2003) (Okamoto et al., 2003), NSR sentinels peaks within minutes and are strongly repressed within 2 to 4 hours following NO_3_^-^ provision (Fig. 2b). As predicted, the sequential deletion of the *HHO* paralogs triggers a de-repression of NSR genes. It is noteworthy that the de-repression of the NSR sentinel genes follows the sequential deletion of HHOs (quite manifest at the 2 and 3 hour time-points) (Fig. 2c). The HRS/HHO quadruple mutant displayed the strongest phenotype and seems to be unable to completely repress the locus even after several hours of NO_3_^-^ provision (Fig. 2b, 2c).

To validate that it is indeed a combination of HRS/HHO deletion that de-repressed the NSR sentinels (as opposed, for example, to the simple effect of *hho3* mutation), we performed a complementation experiment of the quadruple *hrs/hho* mutant with pHRS1:HRS1:GFP. We showed (Fig. 2c) that two independent lines are able to fully restore the WT phenotype regarding the NSR sentinel gene expression. This demonstrates that it is indeed a combination of HRS/HHO deletions that is needed to observe the de-repression of NSR genes (Fig. 2b, 2c, 2d).

These results show that HHOs have a redundant function and that they collectively are involved into repressing NSR sentinels when NO_3_^-^ is provided (Fig. 2).

### HHOs control NO_3_^-^ HATS

Among the three NSR sentinel genes, NRT2.4 and NRT2.5 were shown to be involved in a very high-affinity nitrate transport system (Kiba et al., 2012; Lezhneva et al., 2014). Since we observed interesting molecular phenotypes in the context of N-starvation for the 35S:HRS1 and *hrs1;hho1* mutants (Fig. 1), we tested their high-affinity transport system (HATS) activity following prolonged NO_3_^-^ starvation (Fig. 3a). We performed ^15^NO_3_ labeling experiments, and indeed found that 35S:HRS1 plants are affected in NO_3_^-^ HATS (10 μM) activity. We repeated the experiment to measure the range of affinities at which the HRS1 overexpression had an effect. We observed (Fig. 3b) that the whole high-affinity range was decreased in the 35S:HRS1 plants (with a 2-fold decrease for the very high nitrate affinity conditions, concomitant with a decrease in NRT2.4, NRT2.5, NRT2.1 mRNA accumulation), while low affinity nitrate transport activity remained unchanged in the 35S:HRS1 background (Fig. 3b). The double mutant *hrs1;hho1* displayed little phenotype in this context, as compared to the WT.

**Figure 3.**
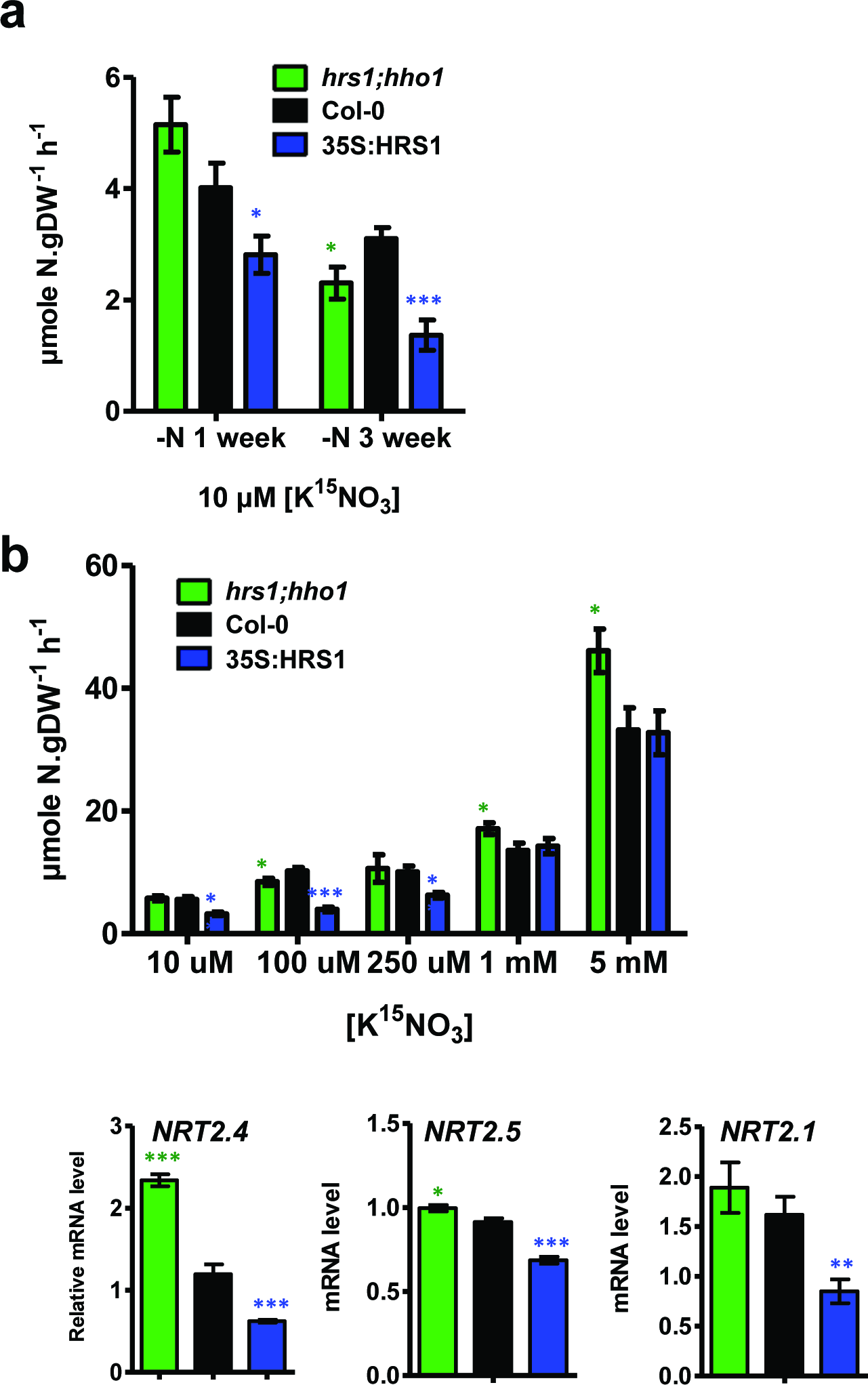
HRS1 and HHO1 negatively control NO_3_^-^ HATS. **(a)** NO_3_^-^ uptake is altered in the 35S:HRS1 and in the double mutant *hrs1;hho1*. Plants were grown for 5 weeks on a N containing media. The media is then shifted to ‐N conditions or +N as a control for 1 or 3 weeks. Values are means ± s.e.m (n= 6). **(b)** One week starved plants were used to quantify NO_3_^-^ HATS and LATS activities as well as high affinity NO_3_^-^ transporter transcript levels. qPCR are normalized to two housekeeping genes (ACT and CLA), values are means ± s.e.m (n= 12). NO_3_ uptake measurements were performed on different ^15^NO_3_ concentrations (10, 100, 250 μM, 1 and 5 mM) to evaluate HATS and LATS. Values are means ± s.e.m (n= 6). The experiment was performed exactly as mentioned for **(a)**. Asterisks indicate significant differences from WT plants (*P<0.05; **P<0.01; ***P<0.001; Student’s t-test).

Since we showed that the *hrs1/hho* mutants were unable to totally repress the *NRT2.4, NRT2.5* loci (Fig. 2b), we also wanted to study functional phenotypes in the HHOs mutants. Thus, we performed ^15^NO_3_ labeling experiment on N-containing media (Fig. 4a). Consistent with our previous findings (Fig. 2a), the sequential deletion of HHOs increases the HATS NO_3_^-^ transport activity with a maximum effect recorded for the *hrs;hho,hho2,hho3* quadruple mutant that displays a 2.5 fold increase of HATS activity (Fig. 4a). Very interestingly, the nitrate transport activity increase is accompanied with a strong stimulation of the quadruple mutant growth in these conditions (Fig. 4b). Although this phenotype is less pronounced than in the quadruple mutant, the double mutant *hrs1;hho1* still displays bigger shoots as compared to the wild type (Sup Fig. 3). The mutant phenotypes were lost in plants grown on ‐N conditions (data not shown). This can be easily explained. Indeed, even if *NRT2.4* and *NRT2.5* are de-repressed in the mutants in both conditions, only the mutants grown on +N can take up more nitrate than WT. Furthermore, HHOs expression is known to be very low in ‐N conditions (Krouk et al., 2010b; Menz et al., 2016), so mutations of genes, being weakly expressed in ‐N, are expected to have low or no effect. These are two possible explanations of why we see mutant phenotypes only on +N conditions.

**Figure 4.**
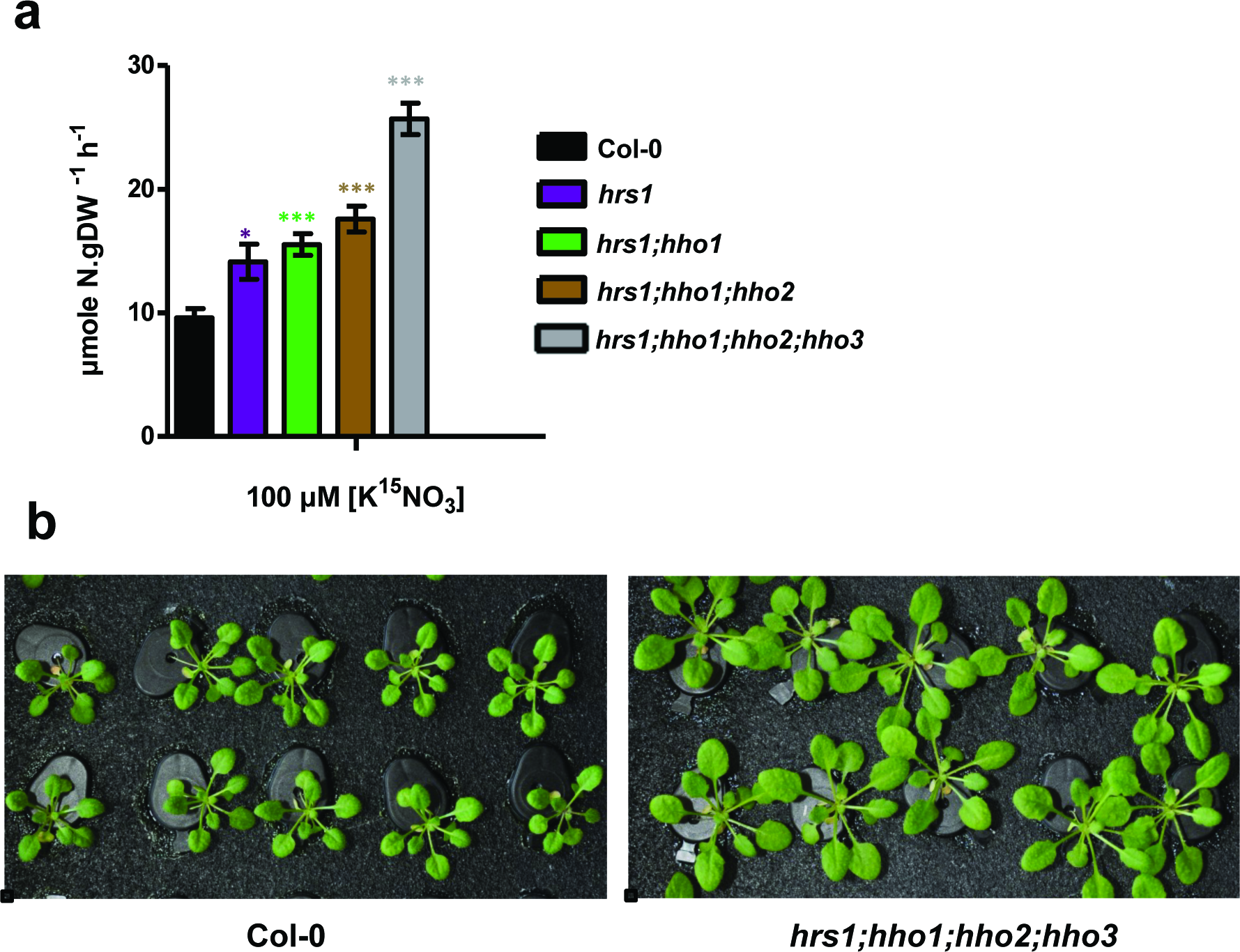
The HHO subfamily represses NO_3_^-^ uptake and growth in +N conditions. **(a)** uptake is altered in the *hrs1, hrs1;hho1, hrs1;hho1;hho2, hrs1;hho1;hho2;hho3* mutants. Plants were grown for 6 weeks on N containing non-sterile hydroponics (0.5 mM NH_4_NO_3_). NO_3_^-^ uptake measurements were performed at 100 μM ^15^NO_3_^-^ to evaluate the HATS. Values are means ± s.e.m (n= 6). Asterisks indicate significant differences from WT plants (*P<0.05; **P<0.01; ***P<0.001; Student’s t-test). **(b)** Representative pictures of Col and the *hrs1;hho1;hho2;hho3* quadruple mutant +N conditions at the day of the uptake experiment show a growth phenotype.

**Figure 5.**
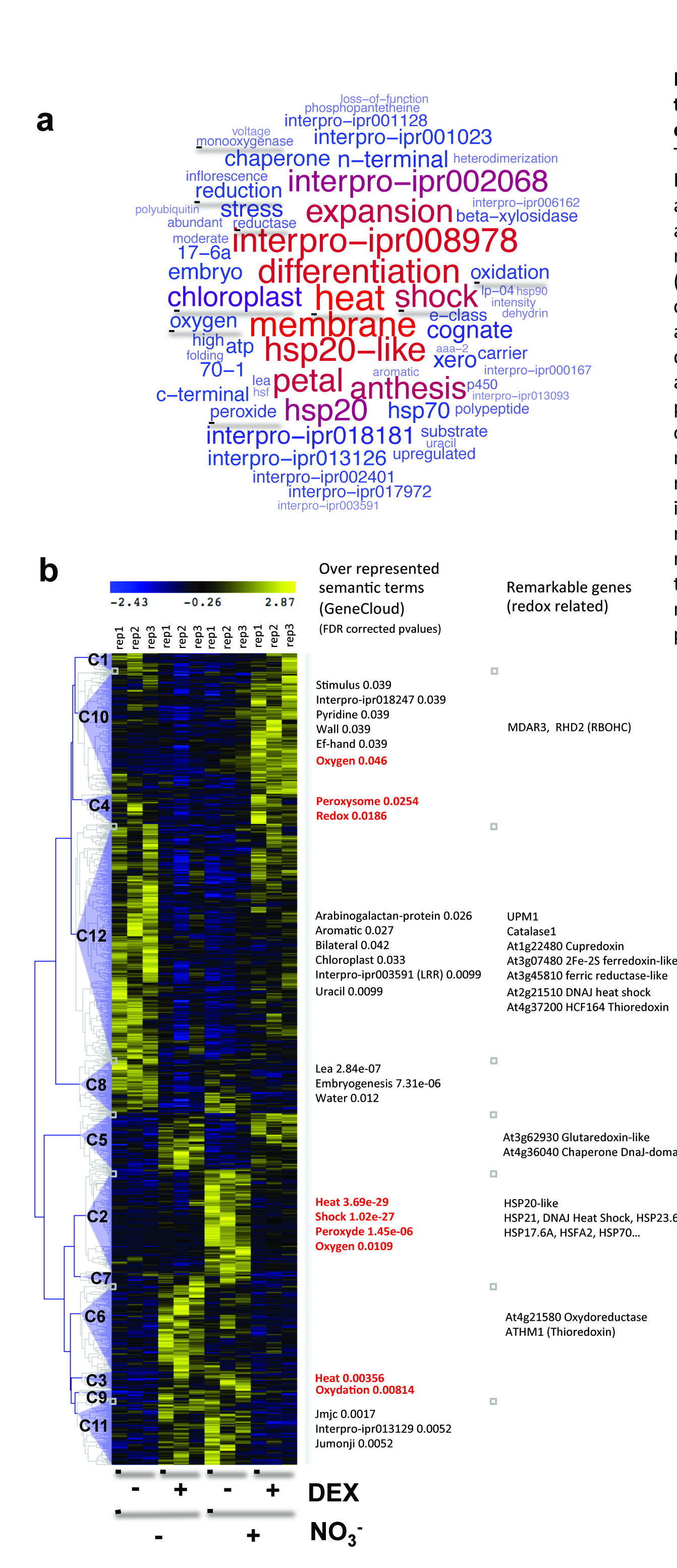
HRS1 direct genome-wide targets are largely NO_3_^-^ dependent and contains many redox related genes. TARGET procedure was performed with NO_3_^-^ (data from Medici et al. (2015)) and without NO_3_^-^ (this work). An ANOVA analysis followed by a Tukey test retrieved 1050 HRS1 regulated genes (ANOVA pval cutoff 0.01, Tukey pval cutoff 0.01, FDR<10%). **(a)** GeneCloud analysis (Krouk et al., 2015) of the 1050 direct targets of HRS1. **(b)** Clustering analysis (Pearson correlation) was performed using MeV software (number of clusters was determined by the FOM method). A selection of over-represented semantic terms is displayed in front of each cluster. Remarkable redox related genes are displayed in the right column. The list of each cluster, their related gene list, as well as their respective semantic analysis are provided in Sup File 1.

In conclusion, the modification of HHOs expression has functional consequences on NO_3_^-^ HATS activities consistent with their role on the transcriptional control on HATS transporters NRT2.4, NRT2.5.

### HRS1 direct targets point to a role of Reactive Oxygen Species

To broaden our investigations around this phenomenon, we studied HRS1 direct genome-wide targets in an N-varying context. To this end, we performed HRS1-perturbations using a *TARGET* analysis (Transient Assay Reporting Genome-wide Effect of Transcription factors (Bargmann et al., 2013; Medici et al., 2015; Para et al., 2014)), including 3 biological repeats, by using root protoplasts not treated with NO_3_^-^ during the *TARGET* procedure. We compared these new results with previously reported *TARGET* results for *HRS1*, in which we maintained NO_3_^-^ in the media (Medici et al., 2015). Both studies where performed in parallel in the exact same conditions and in the same lab. Cycloheximide (CHX) was maintained in the media during the procedure to retrieve only potential direct targets of HRS1. This provided several important insights concerning HRS1 TF activity.

First, we retrieved 1050 potential HRS1 direct targets (non-redundant AGI) (see Material and Method for statistics, and Sup File 1 for the gene list). This list of HRS1 direct targets was subjected to GeneCloud (Krouk et al., 2015) analysis that performs semantic term enrichment analysis on gene lists (Fig. 5a). This revealed that HRS1 controls a highly coherent group of genes, function-wise. Very strikingly, the terms related to redox function (oxidation, peroxide, reductase, oxygen) was highly overrepresented in this list of genes controlled by HRS1 (Fig. 5a, Sup File 1).

A clustering analysis revealed 12 different modes of gene control by HRS1 in combination with NO_3_^-^ (Fig. 5b, cluster lists are provided in Sup File 1). Interestingly, the expressions of the vast majority (86%) of the HRS1 direct genome-wide targets are dependent on the NO_3_^-^ context (Fig. 5b). This can be explained by N related transcriptional, post-transcriptional, post-translational modifications that could affect HRS1 itself or its TF partners.

Among the 12 clusters of HRS1 target genes, the two NO_3_^-^-insensitive ones are Clusters #5 and #8. Cluster #8 contains genes whose functions were reminiscent to the germination control of HRS1 demonstrated in previous work (Wu et al., 2012) (Fig. 5b). Cluster #5 contains genes with diverse function including a Chaperone Dnaj-domain protein and a Glutaredoxin. It is noteworthy that most of the HRS1 regulated gene clusters contain genes related to the redox state of the cells (Figure 5b). However, the most enriched clusters in redox related genes are cluster #12 and #2. Cluster #2 contains many genes annotated as heat shock protein and heat shock factors. For this cluster, HRS1 plays a role of repressor only when NO_3_^-^ is provided to the protoplasts. On the other hand, cluster #12 contains genes that are repressed by HRS1 only when NO_3_^-^ is not present during the TARGET procedure (Fig. 5b). This cluster contains many redox related genes such as a Catalase1 (CAT1), a Ferredoxin, a Thioredoxin, and a Cupredoxin.

In conclusion, this HRS1 TF perturbation analysis in the *TARGET* system prompted us to investigate below i) the role of ROS in the NSR, ii) the role of ROS in the HRS1 dependent control of NSR. It also illustrates that as TF can greatly modify its targets according to a nutritional context (Fig. 5).

### ROS scavenging molecules are strong repressors of NSR

To investigate the role of ROS in NSR, we undertook a pharmacological approach. We showed that NSR is strongly repressed if plants are concomitantly treated with ROS scavenging molecules. In a first experiment, we used a co-treatment with potassium iodide (KI, 5 mM) (Tsukagoshi et al., 2010) and mannitol (5 mM) (Cuin and Shabala, 2007; Shen et al., 1997a; Shen et al., 1997b; Voegele et al., 2005), shown to scavenge H2O2 and HO^•^ respectively. This KI-mannitol co-treatment totally abolished the induction of the NSR sentinel genes (Fig. 6a). In a second experiment, we investigated the respective role of KI and mannitol alone, or in combination and also used diphenyleniodonium (DPI, NADPH oxidases inhibitor) (Orozco-Cardenas et al., 2001; Tsukagoshi et al., 2010) and DMSO as a mock control. This experiment confirmed that the inhibition of ROS production has a severe effect on NSR response. More precisely, for *NRT2.4* and *NRT2.5*, we recorded that KI and DPI strongly repressed the gene response to N deprivation. For *NRT2.5* and *GDH3*, the mannitol alone seems to also have an effect as it dampens down transcriptional responses to N depletion. For *NRT2.5* and *GDH3*, we also found that DMSO may have an effect on its own. However, when DPI treatment was compared to its DMSO mock control, it was significantly affecting NSR for the three sentinel genes. These experiments demonstrate that ROS scavenging molecules are affecting plant NSR. And that ROS are an essential activating potential second messenger of NSR.

**Figure 6.**
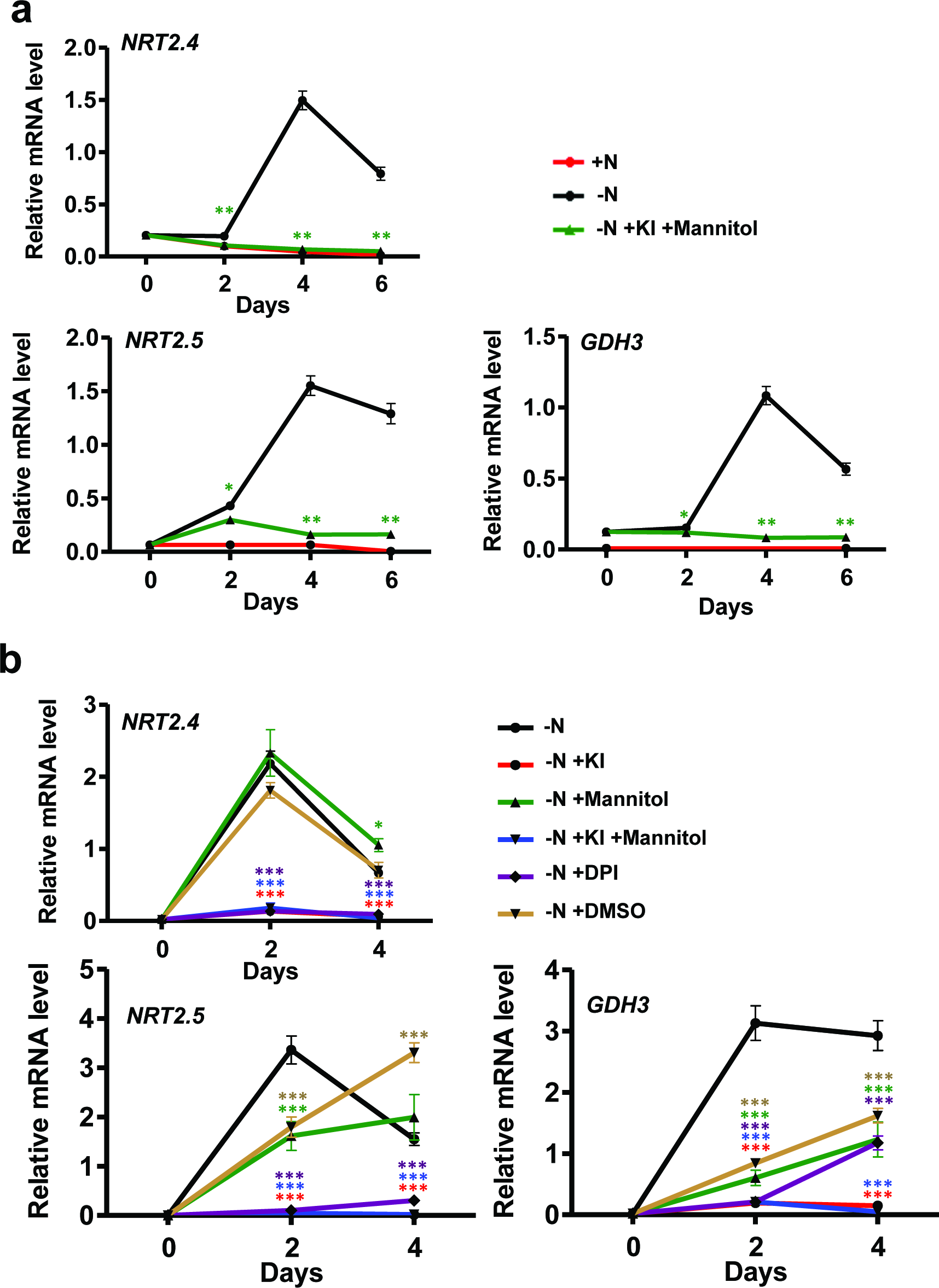
ROS is necessary for the NSR response. **(a)** Altered response of *NRT2.4, NRT2.5* and *GDH3* by KI-mannitol treatment. Plants are grown in sterile hydroponic conditions on N-containing media for 14 days. Thereafter, the media is shifted to ‐N conditions containing or not 5 mM of KI and 5 mM of mannitol for 0, 2, 4, 6 days or +N as a control. **(b)** Altered response of *NRT2.4, NRT2.5* and *GDH3* by ROS scavenger treatment. Plants are grown in sterile hydroponic conditions on N containing media for 14 days. Then plants were transferred to ‐N or +N conditions for 0, 2, 4 days. In parallel, some of the N starved plants were treated with 5 mM KI, 5 mM mannitol, combination of both or with 10 μM DPI. DMSO was used as mock treatment of DPI. All transcript levels were quantified by qPCR and normalized to two housekeeping genes (ACT and CLA), values are means ± s.e.m (n= 4). Asterisks indicate significant differences from WT plants (*P<0.05; **P<0.01; ***P<0.001; Student’s t-test).

Since i) HRS1 regulates ROS related genes (Fig. 5); and because ii) ROS production seem to be important molecules for NSR and; and iii) HRS1 and HHOs are important regulators of NSR themselves (Fig. 1-2); we wanted to know if the NSR control of HRS1 could have some branch being ROS-dependent or, if it was a pure direct regulation. To do this we set up two types of experiments:

-First, we demonstrated that the regulation of NRT2.4 and NRT2.5 loci happens through the potential binding of HRS1 to the promoter of these genes (Fig. 7). To this end, we identified a new potential HRS1 DNA-binding element by running the MEME algorithm (Bailey et al., 2009) on the 500 bp upstream sequences of the most repressed HRS1 genes in the TARGET analysis (List from repressed direct targets in Medici et al. (2015) (Medici et al., 2015)). This new HRS1 cis-element uncovered contains the GANNNTCTNGA consensus that resembles the consensus motif of HHO2 and HHO3 revealed by DAP-seq in the work by O'Malley et al. (2016) (O'Malley et al., 2016) (Fig. 7a). We used EMSA and competition assays with cold probes to demonstrate that HRS1 had a specific affinity for this new motif, and that the conserved cytosine in the sequence is not critical for the DNA-protein recognition, while the distal guanines play a significant role (Fig. 7b-d). Interestingly, this HRS1 motif is found two times in each of the promoters of the 3 sentinel genes as well as in NRT2.1 (Fig. 7e). We thus tested the binding of HRS1 to probes made from the promoter sequences framing the HRS1 binding sites, and validated that HRS1 is able to bind NSR sentinel genes in a promoter context (Fig. 7e-g). Our results are strengthened by DAP-seq data (O'Malley et al., 2016), showing that HHOs sub-family binding (Safi et al., 2017) is specially present in the promoter of the *NRT2.4, NRT2.5, GDH3* genes. Interestingly no specific binding is recorded for KANADI2 or bZIP16 being respectively: a G2-like as well but not in the same sub-family (Safi et al., 2017), or a bZIP (different TF family).

**Figure 7.**
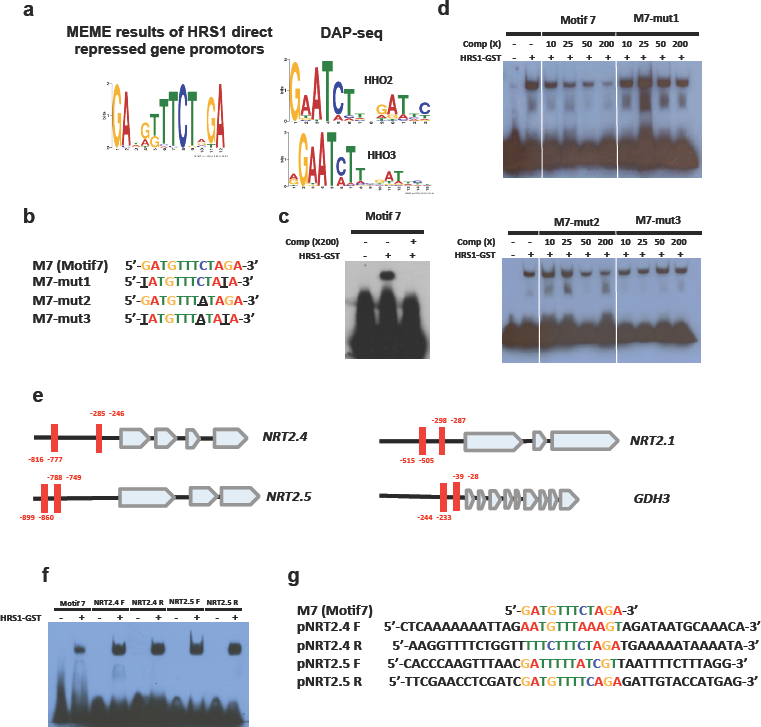
Identification of a new HRS1 c/s-regulatory element and binding of HRS1 to the *NRT2.4* and *NRT2.5* promoters. **(a)** HHOs target motifs. Weight matrix representation of the motifs retrieved by the MEME algorithm analysis from the 500 bp sequences upstream the transcription start sites of the down-regulated HRS1-target genes. HHO2 and HHO3 *cis* motifs retrieved by DAP-seq (O'Malley et al., 2016). **(b)** Wild-type and mutated forms of the HRS1 target motif used in EMSA in c and d. **(c)** EMSA analysis on 48bp (4X repetition of HRS1 target motif); Biotin-TEG labeled DNA probes (20 fmol), GST:HRS1 protein (150 ng). 200X excess of the cold version of the same DNA probes was added in the third well. **(d)** EMSA analysis on 48bp (4X repetition of motifs listed in b); different concentrations of the cold version of the motif or of the 3 mutated forms were added each time. **(e)** Schematic representation of *NRT2.4, NRT2.5, NRT2.1* and *GDH3* genes showing potential HRS1 cis-motif in their promoters. **(f)** EMSA analysis on 40-bp promoter fragments of *NRT2.4* and *NRT2.5*. **(g)** List of the promoter fragments sequences as well as HRS1 target motif used for EMSA analysis. Two promoter fragments sequences were tested for each gene.

Taken together, DAP-seq (Sup Fig. 4), EMSA (Fig. 7), TARGET (Sup Fig. 1a), Overexpression approach (Fig. 1; Sup Fig. 1b), mutant phenotype (Fig. 1-4; Sup Fig. 1b), strongly suggests that HRS1 and its homologous genes directly repress the NSR sentinel genes. Work by Dr Kiba (Riken, Japan) using different techniques also lead to similar conclusions (Kiba et al submitted).

-Second, we wanted to test if the HHO-related phenotypes could also be explained by^®^ a default in a ROS production and/or accumulation. To this aim, by using Amplex Red measurements (Chakraborty et al., 2016; Shin et al., 2005; Shin and Schachtman, 2004), we demonstrated that NSR early (within 6 hours) triggers ROS accumulation in root tissues of WT plants consistent with previous observations (Shin et al., 2005; Shin and Schachtman, 2004). We also found that this accumulation is lost in the 35S:HRS1 plants and reduced in the 35S:HHO1 genotype (Fig. 8a). Unexpectedly, the quadruple hrs/hho mutant did not display any ROS accumulation changes. This could be explained by the fact that HRS1 and HHO1 over-expressors may overproduce ROS scavenging proteins (as found for HRS1 in the TARGET results), when the 4 mutations in the hrs/hho quadruple mutant are not enough to de-repress them at the same level (due to functional redundancy) leading to a WT phenotype for ROS accumulation in the mutant. This interesting observation will be important for future investigations to uncouple the ROS dependent and independent branches of NSR controlled by HRS1 and HHOs.

**Figure 8.**
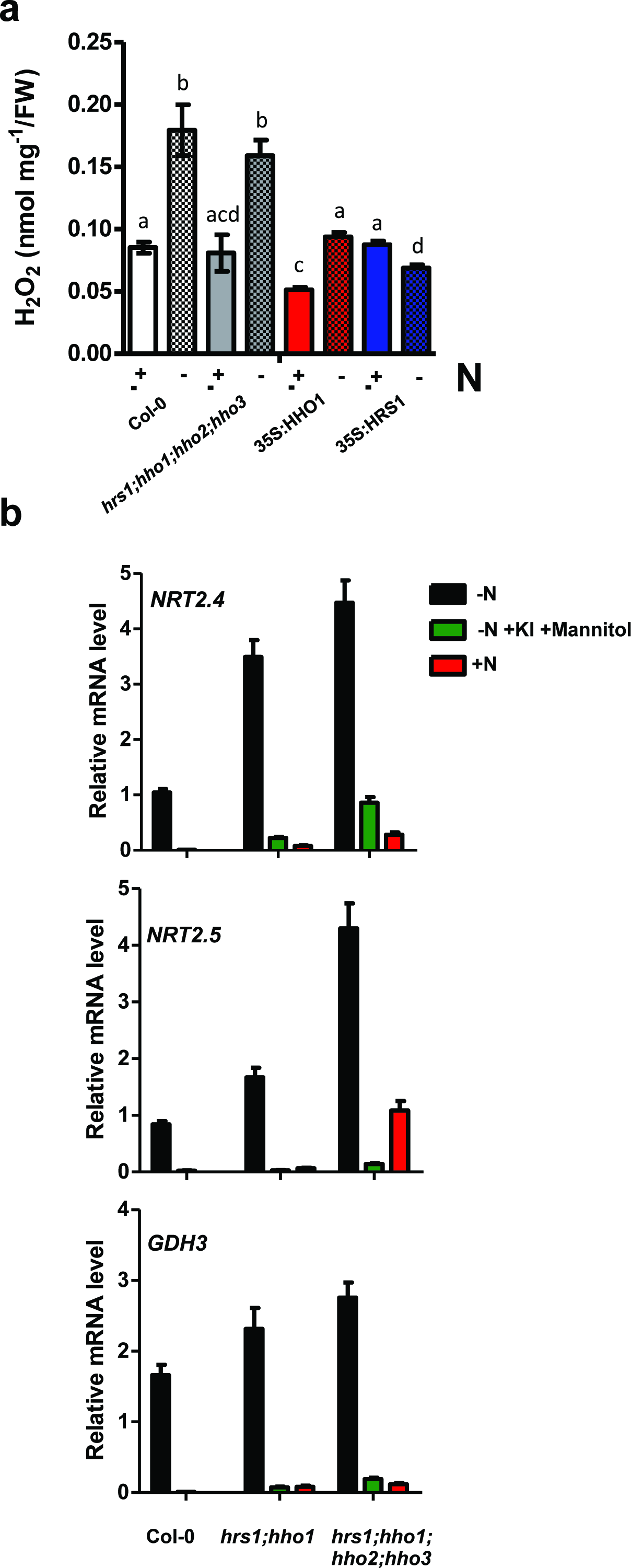
ROS is produced early after nitrogen deprivation, regulated by HRS1 and crucial for the NSR. **(a)** H_2_O_2_ production after N deprivation. Plants were grown in non-sterile hydroponics for 6 weeks on N containing media. Thereafter the media is shifted to ‐N conditions or +N as a control for 6 hours. H_2_O_2_ accumulation was measured using Amplex Red (see Material and Methods). Values are means ± s.e.m (n= 6). Different letters means significantly differences (Student’s t-test, P < 0.05). **(b)** ROS scavenging treatment represses NSR sentinel genes in the *hhos* mutants. Plants are grown in sterile hydroponic conditions on N containing media for 14 days. Plants are then N-deprived for 3 days and treated with 5 mM of KI and 5 mM of mannitol. Plants kept on the same renewed media were used as control. Values are means ± s.e.m (n= 4).

As we observed that ROS early accumulate in response to N starvation (Fig. 8a), we decided to treat the double and quadruple *hrs/hho* mutants with KI-mannitol in this context (Fig. 8b). We found that ROS scavenging treatment indeed represses NSR sentinel genes in the *hhos* mutants to the level of WT, totally abolishing the NSR and the mutant phenotype (Fig. 8b). These results show that ROS scavenging agents are able to overcome the *hhos* mutant phenotype.

In conclusion, we show that HHO genes directly control NSR sentinels (Fig. 7). We also report that upon NSR, ROS are produced and participate to NSR activation. To a certain extent, ROS accumulation can also explain phenotypes observed in HHO affected genotypes. Thus, we conclude that HRS1 and HHO1 are able to reduce ROS accumulation and potentially reduce NSR through a ROS-dependent and parallel branch of the signaling pathway (Fig. 8b and Fig. 9).

**Figure 9.**
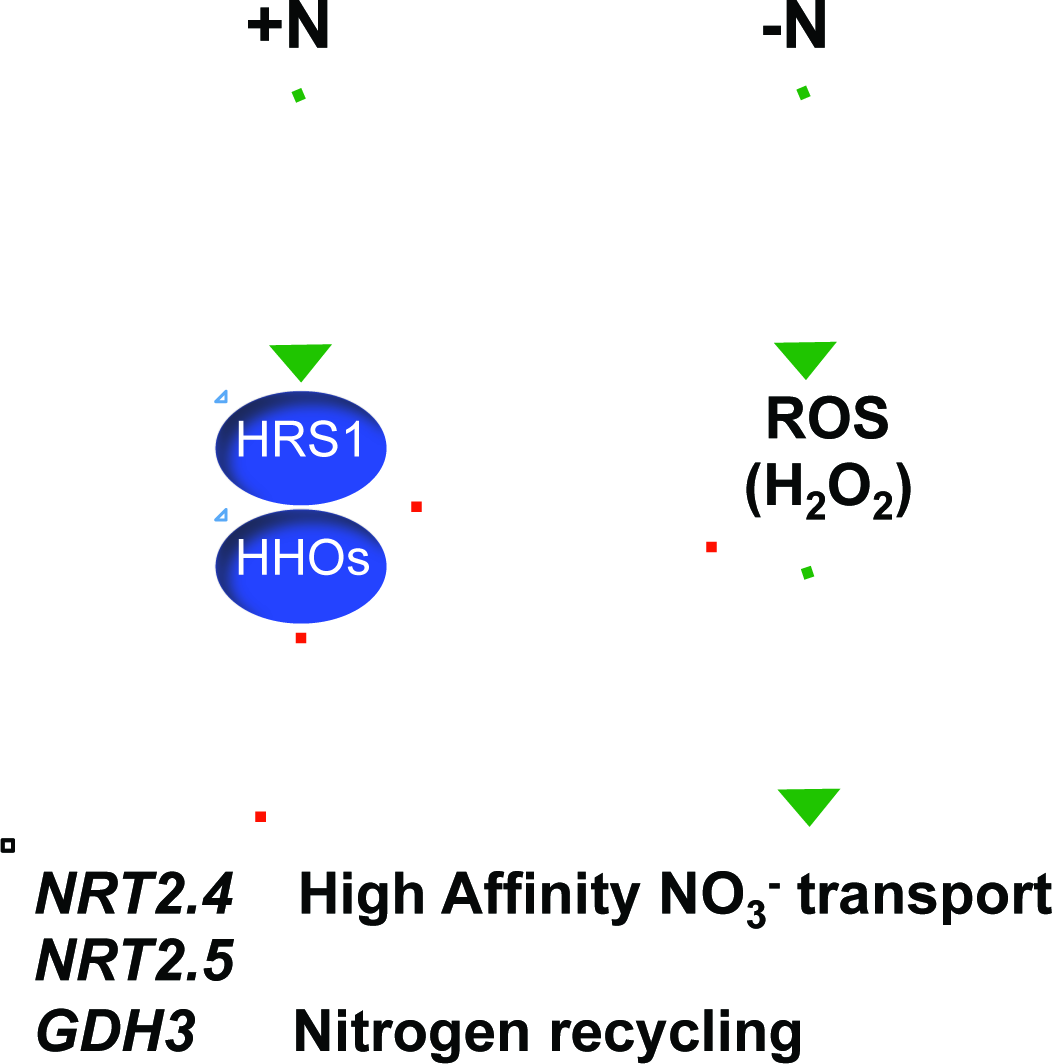
Proposed model of the regulation of NSR by HHOs and ROS. On ‐N conditions, ROS are produced and are needed for the NSR induction. When Nitrogen is present in the media, HRS1 and its homologs are rapidly and highly expressed to repress the NSR either directly by regulating the *NRT2* and *GDH3* promoter activities or indirectly by reducing ROS production.

## Discussion

To date the role of ROS as a potential messenger in NSR was hypothesized by several groups (Krapp et al., 2011; Shin et al., 2005) but, to our knowledge, never formally demonstrated. Previously, Shin et al. (2004) (Shin and Schachtman, 2004) demonstrated that upon +K starvation, ROS accumulation through the action of RHD2 (NADPH-oxidase) was important to sustain the full transcriptional activation of +K transporters. The same group (Shin et al., 2005) also demonstrated that ‐P, and ‐N treatments trigger the production of ROS. However, the direct role of ROS in the NSR was so far elusive. In the current work, we present evidence that, preventing the production of ROS during NSR strongly represses the response of the NSR sentinel genes (Fig. 6). Taken together, these above-mentioned research (Krapp et al., 2011; Shin et al., 2005; Shin and Schachtman, 2004) and others (Balzergue et al., 2017; Hoehenwarter et al., 2016; Mora-Macias et al., 2017; Muller et al., 2015), strongly support that ROS are potential central hubs of the nutrient starvation responses. Because plant are able to differentiate between the different nutritional deprivations, the next challenge will be to understand how is the nutritional specificity of ROS production detected by cells? How is the plant able to detect the ROS signal coming from N rather than a K deficiency? Some clue probably lies into the cellular specificity of the ROS production (Shin et al., 2005). One could also consider that ROS production is an independent and unspecific branch enhancing any kind of nutrient deficiencies which specificities are encoded by genetic factors (as for Phosphate starvation response [PSR], (Puga et al., 2014)). Further research will be necessary to sort this out.

The second factor found in this work, to be a strong regulators of NSR sentinel genes is HRS1. HRS1 was previously found to control the P response of primary root development (Liu et al., 2009; Medici et al., 2015) and to control germination *via* an ABA dependent pathway (Wu et al., 2012). HRS1 and its close HHO homologous genes are also very well known to be among the most NO_3_^-^ regulated gene in the Arabidopsis genome (Canales et al., 2014; Krouk et al., 2010b; Wang et al., 2004). The very strong control of HHOs by NO_3_^-^ seems to be important for its functions. Previously, we showed that HRS1 NO_3_^-^ regulation is necessary to integrate the NO_3_ and the PO_4_^3-^ signal and to trigger appropriate primary root response (Medici et al., 2015). Herein, the show using over-expressors, mutants, TARGET, EMSA, and DAP-seq data, that HRS1 directly represses *NRT2.4, NRT2.5* genes. These genes are activated upon N starvation to retrieve NO_3_^-^ traces in the soil solution. Thus, it seems critical for the plant to repress them (by HRS1 TF activity) when NO_3_^-^ is provided to N starved plants (Fig. 2). In this work, we generated a genotype missing 4 HHOs (HRS1, HHO1, HHO2, HHO3). This genotype is unable to fully repress the NRT2.4 and NRT2.5 loci (Kiba et al., 2012; Lezhneva et al., 2014). These four mutations lead to an important enhancement of the HATS activity (~2.5 fold at 100 μM of external NO_3_^-^), leading to an enhancement of growth (Fig. 4). To our knowledge this opens original perspectives to develop genotypes with increased transport capacities in crops, and improve NUE with potential long-term impact on global warming (see Introduction).

## Conclusion

The model that we propose (Fig. 9) defines two new kinds of molecular actors (HRS1/HHOs and ROS) in the control of plant to NSR (+N → ‐N). We show that ROS are produced upon NSR and that this response production is necessary for NSR sentinel gene activation. We also show that HRS1 and HHO1 i) control ROS accumulation in response to NSR, ii) directly repress NSR sentinel genes (NRT2.4. NRT2.5).

## Material and Methods

### Plant material

The Arabidopsis (*Arabidopsis thaliana*) plants were in the Columbia-0 background. Mutant *hrs1-1* (SALK_067074), *hho1-1* (SAIL_28_D03) (Medici et al., 2015) *hho2-1* (SALK_070096) and *hho3-1* (SALK_146833) were obtained from ABRC seed stock center and homozygote lines were screened by PCR. The double (*hrs1;hho1*), triple (*hrs1;hho1;hho2*) and quadruple (*hrs1;hho1;hho2;hho3*) mutants has been obtained by crossings and PCR validation. The promoter gene *GFP* lines were obtained by PCR and cloning of *HRS1* from genomic sequences, bringing 3-kb upstream promoter region and the gene, into pMDC107 Gateway-compatible vector (Curtis and Grossniklaus, 2003). Overexpressor lines were obtained by cloning HRS1‐ and HHO1-coding sequences into pMDC32 Gateway-compatible vector (see Medici et al., 2015). The constructs were transferred to Agrobacterium tumefaciens strain GV3101 and used for the Arabidopsis transformation by the floral dip method to transform Columbia or the quadruple mutant for complementation (Zhang et al., 2006).

### Growth conditions and treatments

Plants were grown in sterile hydroponic conditions for 14 days in day/night cycles (16/8 h; 90 μmol photons m^-2^s^-1^) at 22°C as described in Krouk et al. (2010) (Krouk et al., 2010b). Hydroponic media consisted of 1 mM KH_2_PO_4_, 1.5 mM MgSO_4_, 0.25 mM K_2_SO_4_, 3 mM CaCl_2_, 0.1 mM Na-Fe-EDTA, 30 μM H_3_BO_3_, 5 μM MnSO_4_, 1 μM ZnSO_4_, 1 μM CuSO_4_, 0.1 μM (NH_4_)_6_Mo_7_O_24_, 5 μM KI; supplied with 3 mM sucrose, 0.5 mM ammonium nitrate (1 mM KNO_3_ for results in figure 1) and 0.5 g L^-1^ MES. pH was adjusted to 5.7 by adding KOH 1M. Plants were grown for 14 days in day/night cycles (16/8 h; 90 μmol photons m^-2^.s^-1^) at 22 °C. For +N → ‐N experiments, plants were transferred to an equivalent fresh nitrogen-free medium or 0.5 mM ammonium nitrate containing medium (1 mM KNO_3_ for results in figure 1). For ‐N → +N experiments, all plants have been nitrogen starved for 3 days, and then transferred to an equivalent fresh medium containing 0.5 mM ammonium nitrate. Drug treatments for ROS scavenging were added upon plants transfer to the new media. Roots (corresponding to approximately 60 plants coming from a single phytatray) were harvested at different time points and immediately frozen in liquid nitrogen. Each time point and genotype being harvested from a different phytatray. Experiments have been performed at least twice and representative data are reported in figures.

For non-sterile hydroponic culture, seeds were sown on Eppendorf taps with 1 mm whole filled by H_2_O-agar 0.5% solution and grown during 7 days on H_2_O. Then, plants were transferred to the same media as above (without sucrose) and supplied with 0.5 mM ammonium nitrate and grown in short day light period (8h light 23°C and 16h dark 21°C) at 260 μmol photons m^-2^.s^-1^ and 70% humidity. Nutritive solution was renewed every 4 days for 5 weeks. Then plants were transferred to an equivalent fresh nitrogen-free medium or 0.5 mM ammonium nitrate containing medium (1 to 3 weeks for ^15^NO_3_^-1^ uptake experiments and 6h for ROS measurements).

For mutants complementation experiments (Fig. 2c), Columbia and *hrs1;hho1;hho2;hho3;pHRS1:HRS1:GFP* sterilized seeds were sown on the surface of sterile solid media (1% (w/v) agar) consisting of MS/2 basal salt medium containing no nitrogen, supplemented with 3 mM sucrose, 0.5 mM ammonium nitrate, MES buffered at pH 5.7 (0.5 g L^-1^). Agar plates were incubated vertically in *in vitro* growth chamber for 12 days in day/night cycles (16/8 h; 90 μmol photons m^-2^.s^-1^) at 22 °C.

### Real-time qPCR analysis

Total RNAs were extracted from Arabidopsis roots using Trizol^®^ and digested with DNAseI (Sigma-Aldrich, St Louis, USA). RNAs were then reverse transcribed to one-strand complementary DNA using Thermo script RT (Invitrogen) according to the manufacturer’s protocol. Gene expression was determined by quantitative PCR (LightCycler 480; Roche Diagnostics, Basel, Switzerland) using gene-specific primers (provided upon request) and TAKARA mix (Roche, IN, USA). Expression levels of tested genes were normalized to expression levels of the actin and clathrin genes as previously described in (Krouk et al., 2010b).

### Expression and purification of GST-HRS1 protein

The protocol was fully described in (Medici et al., 2015). Briefly, *HRS1* coding sequence were first cloned in the pDONR207 vector, and then transferred to pDEST15 vector (Invitrogen) by LR reaction following the manufacturer’s instructions. The GST-HRS1 fusion protein was expressed in Escherichia coli Rosetta 2(DE3)pLysS (Novagen, Darmstadt, Germany). Transformed cells were grown in a phosphate-buffered rich medium (Terrific broth) at 37°C containing appropriate antibiotics until the OD660 reached 0.7-0.8. After induction with 1 mM IPTG (isopropyl-b-D-thiogalactoside) for 16 h at 22° C, bacteria were harvested by centrifugation (6000 xg, 10 min, 4°C) and suspended in 1× PBS buffer containing lysozyme from chicken egg white (Sigma) and complete protease inhibitor cocktail (Roche). The resulting cell suspension was sonicated and centrifuged at 15,000 g, for 15 min at 4°C to remove intact cells and debris. The proteins extract was mixed with buffered glutathione sepharose beads (GE Healthcare, Freiburg, Germany), and incubated at 4°C for 3 h. The resin was centrifuged (500 xg, 10 min, 4°C) and washed five times with 1× PBS buffer.

GST-HRS1 was then eluted with 10 mM reduced glutathione (Sigma) in 50 mM Tris buffer and dialyzed overnight in 150 mM NaCl, 50 mM HEPES, pH 7.4 buffer.

All fractions were subjected to SDS-PAGE, and proteins concentrations were determined. For proteins quantification, absorbance measurements were recorded on a nanodrop spectrophotometer (Model No.1000, Thermo Scientific Inc., Wilmington, Delaware, USA) at 280 nm, and in parallel on a VICTOR2™ microplate reader (MULTILABEL COUNTER, life sciences) at 660 nm using the Pierce 660 nm Protein Assay (Pierce/Thermo Scientific, Rockford)

### Electophoretic Mobility Shift Assay

EMSA was performed using GST-HRS1 purified protein and DNA probes labeled with Biotin-TEG at the 3′ end. Biotin-TEG 3′ end ‐labeled single-stranded DNA oligonucleotides were incubated at 95 °C for 10 min and then annealed to generate double-stranded DNA probes by slow cooling. The sequences of the oligonucleotide probes were synthesized by Eurofins Genomics and are provided in Figure 7. The binding of the purified proteins (~ 150 ng) to the Biotin-TEG labeled probes (20 fmol) was carried out using the LightShift Chemiluminescent EMSA Kit (Thermo Scientific, Waltham, USA) in 20 μL reaction mixture containing 1× binding buffer (10 mM Tris, 50 mM KCl, 1 mM DTT, pH 7.5), 2.5% glycerol, 5 mM MgCl2, 2 μg of poly (dI-dC) and 0.05 % NP-40. After incubation at 24° C for 30 min, the protein-probe mixture was separated in a 4% polyacrylamide native gel at 100 V for 50 min then transferred to a Biodyne B Nylon membrane (Thermo Scientific) by capillary action in 20X SSC buffer overnight. After ultraviolet crosslinking (254 nm) for 90 s at 120 mJ.cm^-2^, the migration of Biotin-TEG labeled probes was detected using horseradish peroxidase-conjugated streptavidin in the LightShift Chemiluminescent EMSA Kit (Thermo Scientific) according to the manufacturer's protocol, and then exposed to X-ray film.

### ROS measurement

An Amplex^®^ Red Hydrogen Peroxide/Peroxidase Assay Kit (Molecular probes, Eugene, OR, USA) was used to measure H_2_O_2_ production in 6-week-old plants, according to the manufacturer’s protocol. Six independent roots for each treatment and genotype were frozen and ground in liquid N. All extraction protocol was carried out in a cold room at 4 °C. Two hundred μl of phosphate buffer (50 mM K_2_HPO_4_, pH 7.4) was added to 50 mg of ground frozen tissue. After 2 centrifugations at 14000 g for 10 min, 50 μl of the supernatant was added to 50 μl of the Amplex^®^ Red mixture (100 μM (10-acethyl-3,7-dihydrophenoxazine) and 0.2 U/ml horseradish peroxidase) at room temperature (25 °C) for 30 min under dark conditions. The absorbance was measured using VICTOR2^™^ microplate reader (MULTILABEL COUNTER, life sciences) at 560 nm in 96-well transparent plates. A blank (containing 50 μL phosphate buffer and 50 μL Amplex^®^ Red reagent) was considered as a negative control. H2O2 quantities were reported to the exact powder mass of each sample. The absorbance was measured twice in each point.

### ^15^NO_3_^-^ uptake

Influx of ^15^NO_3_^-^ into the roots was assayed as described previously (Lejay et al., 1999). The plants were sequentially transferred to 0.1 mM CaSO_4_ for 1 min and then, to basal nutrient medium (pH 5.7) containing appropriate concentrations of K^15^NO_3_. In the labeling solution, ^15^NO_3_^-^ was used at 10 to 250 μM for HATS and 1 to 5 mM for LATS. After 5 min, roots were washed for 1 min in 0.1 mM CaSO4, harvested, dried at 70°C for 48 h and analyzed. The total N content and atomic percentage ^15^N abundance of the samples were determined by continuous-flow mass spectrometry, as described previously (Clarkson et al., 1996), using a Euro-EA Eurovector elemental analyzer coupled with an IsoPrime mass spectrometer (GV Instruments). Each uptake value is the mean +/‐ SE of 6 replicates (6 independent roots from different plants).

### Phylogenetic analysis

The phylogeny reconstruction was established as described in (Safi et al., 2017) on the whole protein sequences. Briefly, the tree was built using the mafft algorithm (http://mafft.cbrc.jp/alignment/server/, parameters: G-INS-1, BLOSUM62) and drawn with FigTree. Different parameters including FFT-NS-2, FFT-NS-i, E-INS-i were used that yielded very similar trees.

### TF perturbation assays in the TARGET system

The TARGET procedure has been performed as previously described in and (Bargmann et al., 2013; Medici et al., 2015). Protoplasts were treated with 35 μM CHX (Cycloheximide) for 30 min, then 10 μM DEX (Dexamethasone) was added and cell suspension was incubated in the dark over night at room temperature. Controls were respectively treated with DMSO (dimethylsulfoxide) and ethanol.

The red fluorescent protein was used as marker selection for fluorescent-activated cell sorting of successfully transformed protoplasts. Free nitrogen buffer was maintained during the whole procedure (as compared to Medici et al. (2015) during which NO_3_^-^ was maintained during the TARGET procedure). RNA was extracted and amplified for hybridization with ATH1 Affymetrix chips.

Transcriptomic analysis was performed using ANOVA followed by a Tukey test using R (https://www.r-project.org/) custom made scripts following previously published procedures (Obertello et al., 2010; Ristova et al., 2016). Clustering was perfomed using the MeV software (http://mev.tm4.org/).

Briefly, the ANOVA analysis was carried out using the R *aov*() function on log2 MAS5-normalized data. A probe signal has been modeled as follows: Yi = α_1_.DEX + α_2_.NO_3_ + α_3_.NO3*DEX + ε; where α_1_ to α_3_ represent the coefficient quantifying the effect of each of the factors (DEX, NO_3_) and their interaction (DEX*NO_3_), and ε represents the non-explained variance. We determined the false discovery rate (FDR) to be <10% for an ANOVA pvalue-cutoff of 0.01 and a Tuckey pvalue-cutoff of 0.01.

## Acknowledgements

This work was supported by Agence Nationale de la Recherche (IMANA ANR-14-CE19-0008 with a doctoral fellowship to AS), by CNRS (LIA-CoopNet, PEPS-SuperRegNet) and by NSF (IOS 1339362-NutriNet).

---

**Supplementary Figure 1.**
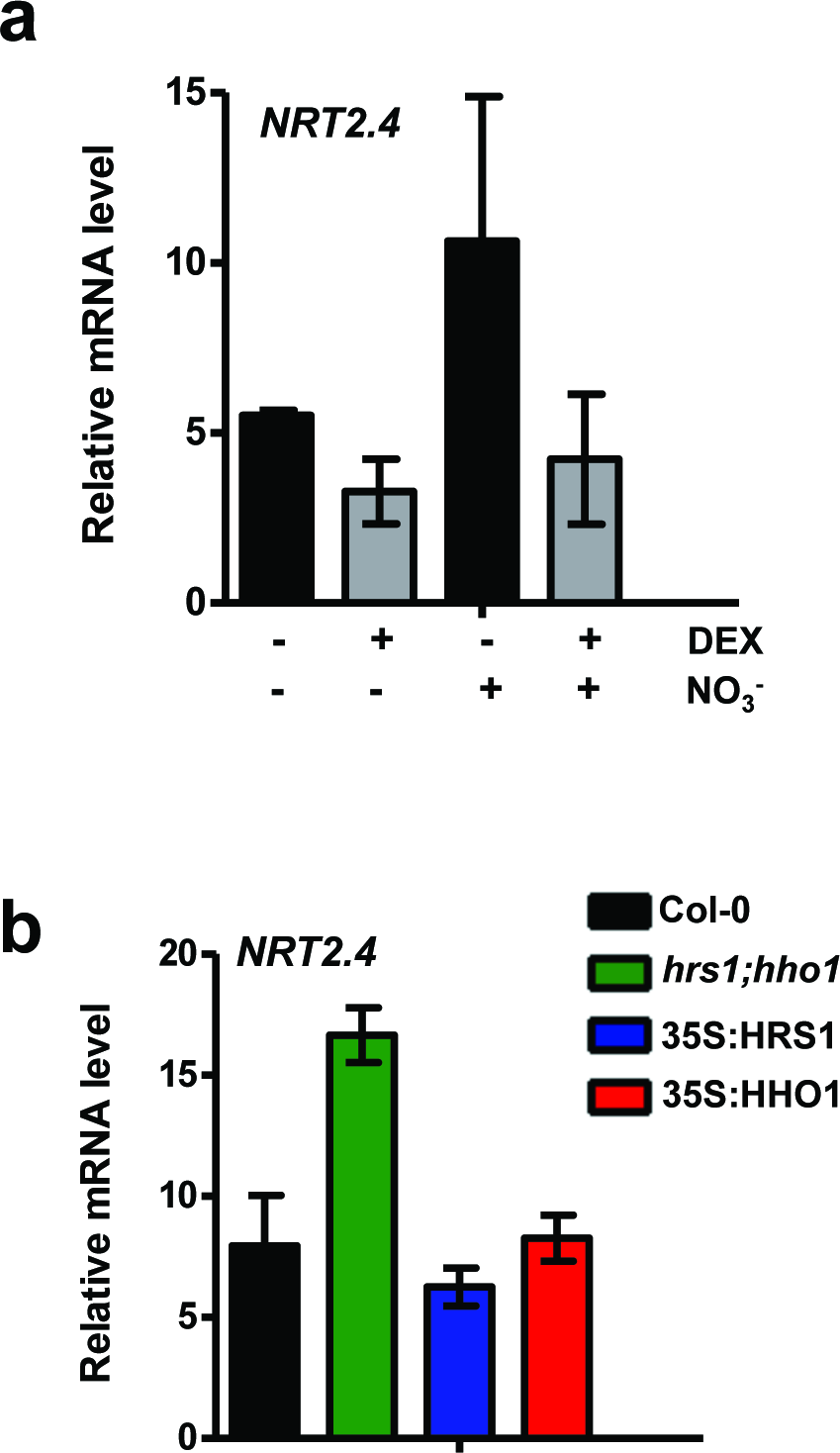
*NRT2.4* is down-regulated by HRS1 and HHO1. **(a)** HRS1 TARGET shows a direct repression of NRT2.4 in Arabidopsis root protoplasts data from Medici et al., (2015) **(b)** *NRT2.4* transcripts level in Wild-type, *hrs1;hho1*, 35SHRS1 and HHO1 genotypes (data from Medici et al., 2015).

**Supplementary Figure 2.**
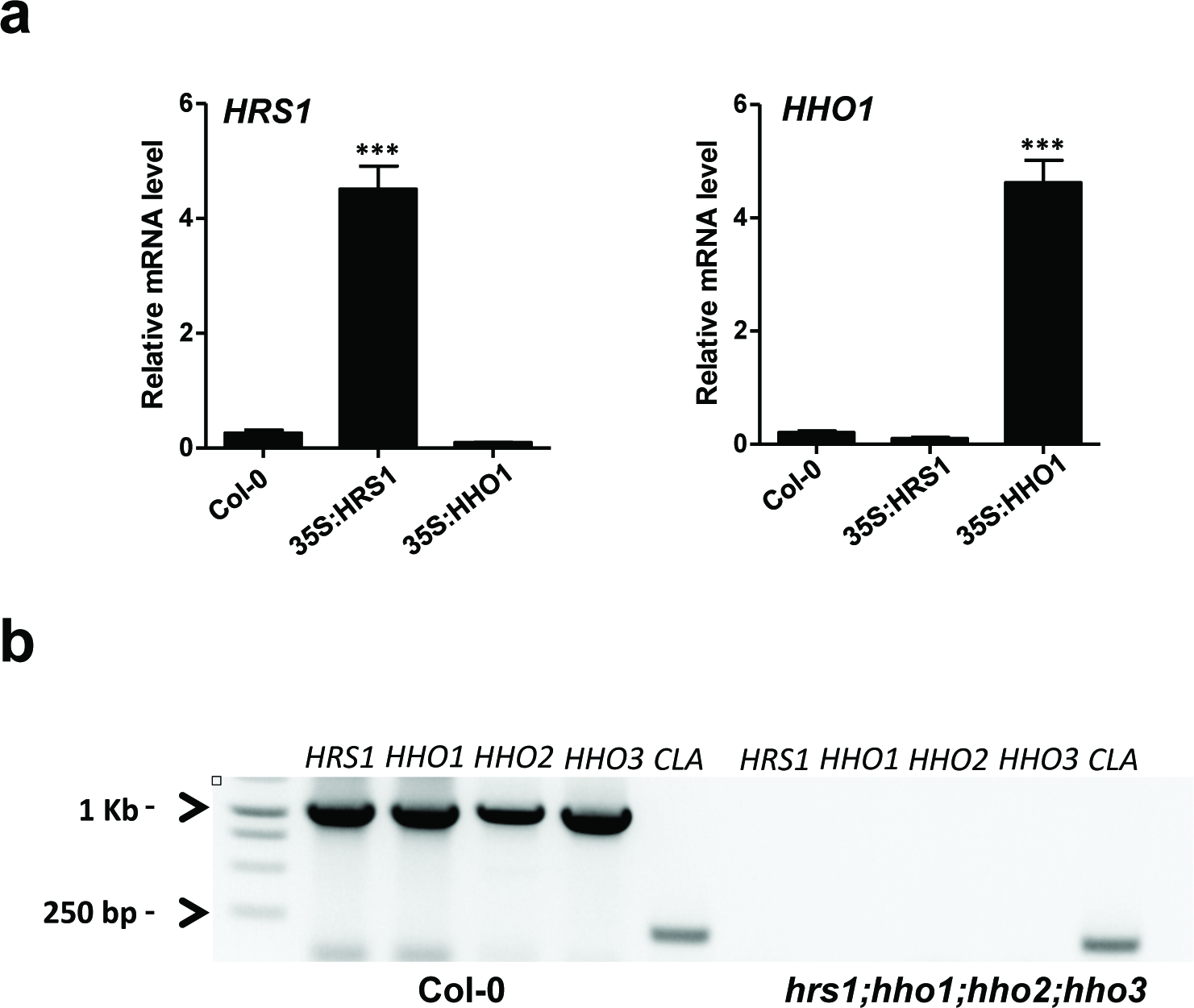
Characterization of quadruple mutant and OE lines. **(a)** Characterization of HRS1 and HHO1 mRNA relative accumulations in 35S:HRS1 and 35S:HHO1 transgenic plants. **(b)** Characterization of the *hrs1/hho* quadruple mutant. The absence of full length transcripts for HRS1, HHO1, HHO2 and HHO3 was verified by RT-PCR (PCR ATG-Stop) on mRNA isolated from roots of *hrs1;hho1;hho2;hho3* mutant grown for 14 days on MS/2 media (Clathrin was used as control).

**Supplementary Figure 3.**
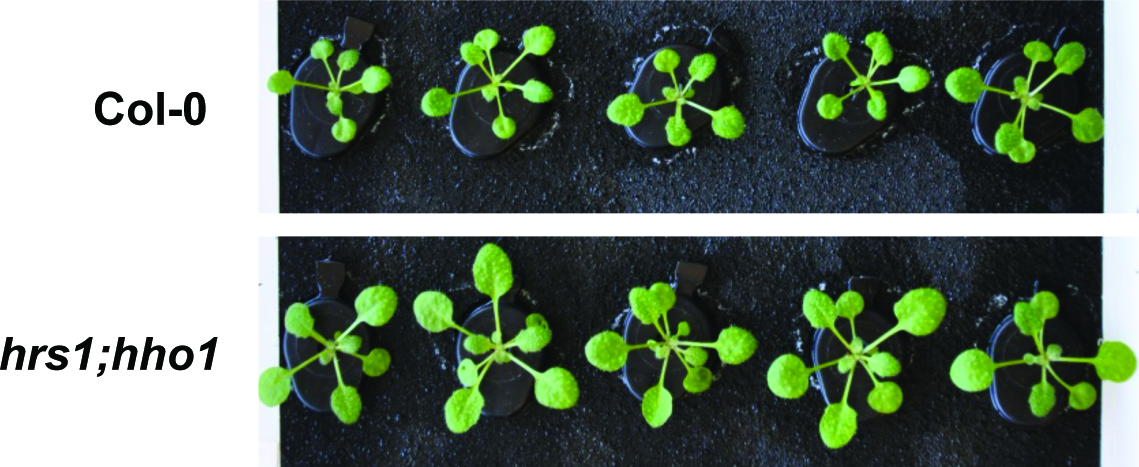
*hrs1;hho1* double mutant displays growth phenotype in +N conditions. Representative pictures of Col and the *hrs1;hho1* double mutant. Plants were grown for 6 weeks on N containing non-sterile hydroponics (0.5 mM NH4NO_3_).

**Supplementary Figure 4.**
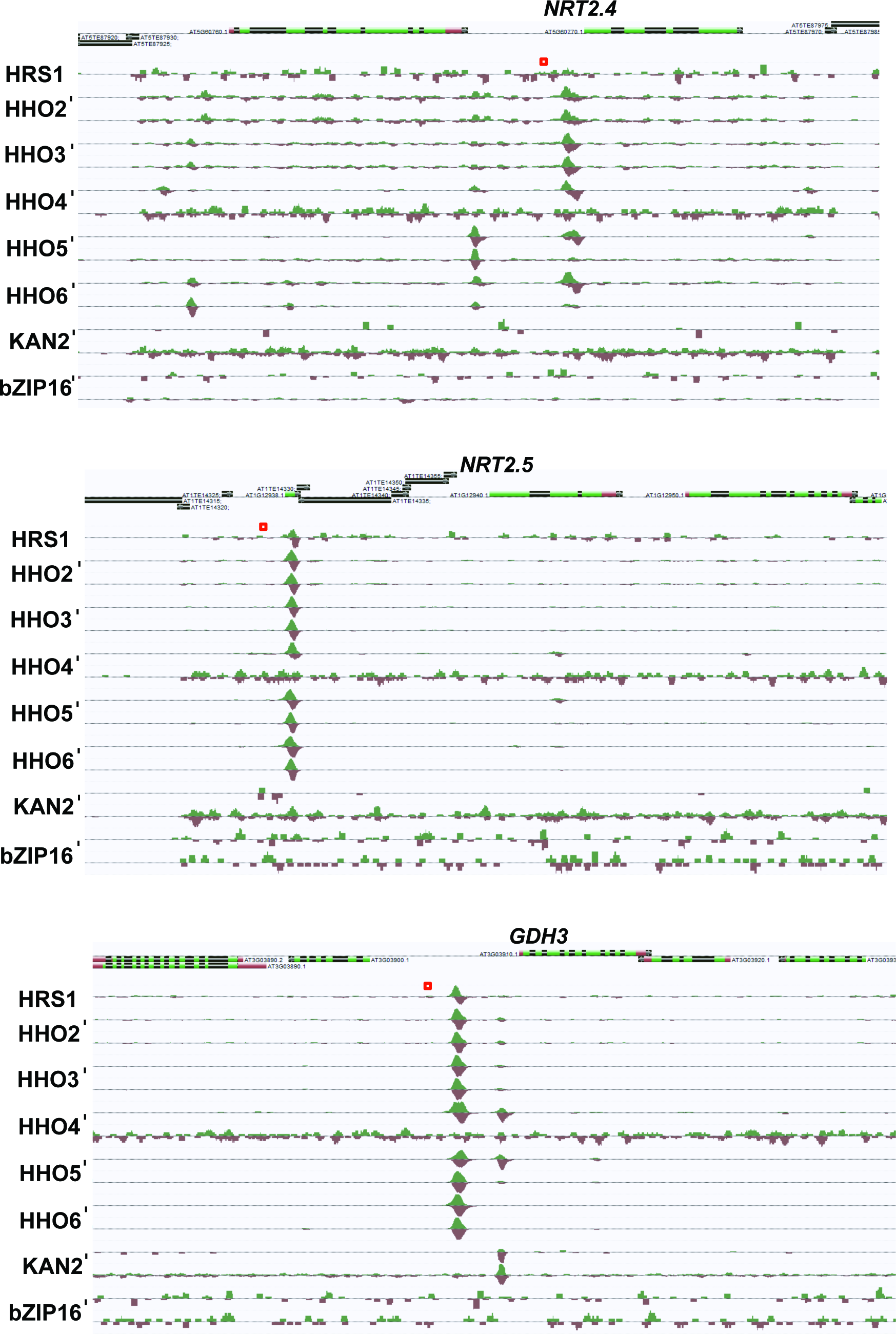
HHOs binding evidences in *NR2.4, NRT2.5* and *GDH3* promoters. Data from DAP-seq (O'Malley et al., 2016) showing binding peaks of HHO family TFs in NR2.4, NRT2.5 and GDH3 promoters. KAN2 (GARP family) and bZIP16 binding profiles were used as controls. The second line of each TF correspond to the demethylated DNA version (ampDAP-seq) used for DAP-seq binding.

## References

Alboresi, A., Gestin, C., Leydecker, M.T., Bedu, M., Meyer, C., and Truong, H.N. (2005). Nitrate, a signal relieving seed dormancy in Arabidopsis. Plant Cell Environ. 28:500–512.

Alvarez, J.M., Riveras, E., Vidal, E.A., Gras, D.E., Contreras-Lopez, O., Tamayo, K.P., Aceituno, F., Gomez, I., Ruffel, S., Lejay, L., et al. (2014). Systems approach identifies TGA1 and TGA4 transcription factors as important regulatory components of the nitrate response of Arabidopsis thaliana roots. Plant J 80:1–13.

Araus, V., Vidal, E.A., Puelma, T., Alamos, S., Mieulet, D., Guiderdoni, E., and Gutierrez, R.A. (2016). Members of BTB Gene Family of Scaffold Proteins Suppress Nitrate Uptake and Nitrogen Use Efficiency. Plant Physiol 171:1523–1532.

Bailey, T.L., Boden, M., Buske, F.A., Frith, M., Grant, C.E., Clementi, L., Ren, J., Li, W.W., and Noble, W.S. (2009). MEME SUITE: tools for motif discovery and searching. Nucleic Acids Res 37:W202–208.

Balzergue, C., Dartevelle, T., Godon, C., Laugier, E., Meisrimler, C., Teulon, J.M., Creff, A., Bissler, M., Brouchoud, C., Hagege, A., et al. (2017). Low phosphate activates STOP1-ALMT1 to rapidly inhibit root cell elongation. Nat Commun 8:15300.

Bargmann, B.O., Marshall-Colon, A., Efroni, I., Ruffel, S., Birnbaum, K.D., Coruzzi, G.M., and Krouk, G. (2013). TARGET: a transient transformation system for genome-wide transcription factor target discovery. Mol Plant 6:978–980.

Canales, J., Moyano, T.C., Villarroel, E., and Gutierrez, R.A. (2014). Systems analysis of transcriptome data provides new hypotheses about Arabidopsis root response to nitrate treatments. Front Plant Sci 5:22.

Castaings, L., Camargo, A., Pocholle, D., Gaudon, V., Texier, Y., Boutet-Mercey, S., Taconnat, L., Renou, J.P., Daniel-Vedele, F., Fernandez, E., et al. (2009). The nodule inception-like protein 7 modulates nitrate sensing and metabolism in Arabidopsis. Plant J 57:426–435.

Castro Marin, I., Loef, I., Bartetzko, L., Searle, I., Coupland, G., Stitt, M., and Osuna, D. (2010). Nitrate regulates floral induction in Arabidopsis, acting independently of light, gibberellin and autonomous pathways. Planta.

Chakraborty, S., Hill, A.L., Shirsekar, G., Afzal, A.J., Wang, G.L., Mackey, D., and Bonello, P. (2016). Quantification of hydrogen peroxide in plant tissues using Amplex Red. Methods 109:105–113.

Clarkson, D.T., Gojon, A., Saker, L.R., Wiersema, P.K., Purves, J.V., Tillard, P., Arnold, G.M., Paams, A.J.M., Waalburg, W., and Stulen, I. (1996). Nitrate and ammonium influxes in soybean (Glycine max) roots: Direct comparison of ^13^N and ^15^N tracing. Plant Cell Environ. 19:859–868.

Crawford, N.M., and Glass, A.D.M. (1998). Molecular and physiological aspects of nitrate uptake in plants. Trends in Plant Science 3:389–395.

Cuin, T.A., and Shabala, S. (2007). Compatible solutes reduce ROS-induced potassium efflux in Arabidopsis roots. Plant Cell Environ 30:875–885.

Curtis, M.D., and Grossniklaus, U. (2003). A gateway cloning vector set for high-throughput functional analysis of genes in planta. Plant Physiol 133:462–469.

Forde, B.G., and Walch-Liu, P. (2009). Nitrate and glutamate as environmental cues for behavioural responses in plant roots. Plant Cell Environ 32:682–693.

Gansel, X., Munos, S., Tillard, P., and Gojon, A. (2001). Differential regulation of the NO_3_^-^ and NH_4_+ transporter genes AtNrt2.1 and AtAmt1.1 in Arabidopsis: relation with longdistance and local controls by N status of the plant. Plant J 26:143–155.

Gruber, B.D., Giehl, R.F., Friedel, S., and von Wiren, N. (2013). Plasticity of the Arabidopsis root system under nutrient deficiencies. Plant Physiol 163:161–179.

Guan, P., Ripoll, J.J., Wang, R., Vuong, L., Bailey-Steinitz, L.J., Ye, D., and Crawford, N.M. (2017). Interacting TCP and NLP transcription factors control plant responses to nitrate availability. Proc Natl Acad Sci U S A 114:2419–2424.

Han, M., Okamoto, M., Beatty, P.H., Rothstein, S.J., and Good, A.G. (2015). The Genetics of Nitrogen Use Efficiency in Crop Plants. Annu Rev Genet 49:269–289.

Ho, C.H., Lin, S.H., Hu, H.C., and Tsay, Y.F. (2009). CHL1 functions as a nitrate sensor in plants. Cell 138:1184–1194.

Hoehenwarter, W., Monchgesang, S., Neumann, S., Majovsky, P., Abel, S., and Muller, J. (2016). Comparative expression profiling reveals a role of the root apoplast in local phosphate response. bMc Plant Biol 16:106.

Hu, H.C., Wang, Y.Y., and Tsay, Y.F. (2009). AtCIPK8, a CBL-interacting protein kinase, regulates the low-affinity phase of the primary nitrate response. Plant J 57:264–278.

Kiba, T., Feria-Bourrellier, A.B., Lafouge, F., Lezhneva, L., Boutet-Mercey, S., Orsel, M., Brehaut, V., Miller, A., Daniel-Vedele, F., Sakakibara, H., et al. (2012). The Arabidopsis nitrate transporter NRT2.4 plays a double role in roots and shoots of nitrogen-starved plants. Plant Cell 24:245–258.

Kiba, T., and Krapp, A. (2016). Plant Nitrogen Acquisition Under Low Availability: Regulation of Uptake and Root Architecture. Plant Cell Physiol 57:707–714.

Kotur, Z., and Glass, A.D. (2015). A 150 kDa plasma membrane complex of AtNRT2.5 and AtNAR2.1 is the major contributor to constitutive high-affinity nitrate influx in Arabidopsis thaliana. Plant Cell Environ 38:1490–1502.

Krapp, A., Berthome, R., Orsel, M., Mercey-Boutet, S., Yu, A., Castaings, L., Elftieh, S., Major, H., Renou, J.P., and Daniel-Vedele, F. (2011). Arabidopsis roots and shoots show distinct temporal adaptation patterns toward nitrogen starvation. Plant Physiol 157:1255–1282.

Krouk, G. (2017). Nitrate signalling: Calcium bridges the nitrate gap. Nat Plants 3:17095.

Krouk, G., Carre, C., Fizames, C., Gojon, A., Ruffel, S., and Lacombe, B. (2015). GeneCloud Reveals Semantic Enrichment in Lists of Gene Descriptions. Mol Plant 8:971–973.

Krouk, G., Crawford, N.M., Coruzzi, G.M., and Tsay, Y.F. (2010a). Nitrate signaling: adaptation to fluctuating environments. Curr Opin Plant Biol 13:266–273.

Krouk, G., Mirowski, P., LeCun, Y., Shasha, D.E., and Coruzzi, G.M. (2010b). Predictive network modeling of the high-resolution dynamic plant transcriptome in response to nitrate. Genome Biol 11:R123.

Lejay, L., Tillard, P., Lepetit, M., Olive, F., Filleur, S., Daniel-Vedele, F., and Gojon, A. (1999). Molecular and functional regulation of two NO_3_^-^ uptake systems by N‐ and C-status of Arabidopsis plants. Plant J 18:509–519.

Leran, S., Edel, K.H., Pervent, M., Hashimoto, K., Corratge-Faillie, C., Offenborn, J.N., Tillard, P., Gojon, A., Kudla, J., and Lacombe, B. (2015). Nitrate sensing and uptake in Arabidopsis are enhanced by ABI2, a phosphatase inactivated by the stress hormone abscisic acid. Sci Signal 8:ra43.

Lezhneva, L., Kiba, T., Feria-Bourrellier, A.B., Lafouge, F., Boutet-Mercey, S., Zoufan, P., Sakakibara, H., Daniel-Vedele, F., and Krapp, A. (2014). The Arabidopsis nitrate transporter NRT2.5 plays a role in nitrate acquisition and remobilization in nitrogen-starved plants. Plant J 80:230–241.

Li, Y., Krouk, G., Coruzzi, G.M., and Ruffel, S. (2014). Finding a nitrogen niche: a systems integration of local and systemic nitrogen signalling in plants. J Exp Bot.

Liu, H., Yang, H., Wu, C., Feng, J., Liu, X., Qin, H., and Wang, D. (2009). Overexpressing HRS1 confers hypersensitivity to low phosphate-elicited inhibition of primary root growth in Arabidopsis thaliana. J Integr Plant Biol 51:382–392.

Liu, K.H., Niu, Y., Konishi, M., Wu, Y., Du, H., Sun Chung, H., Li, L., Boudsocq, M., McCormack, M., Maekawa, S., et al. (2017). Discovery of nitrate-CPK-NLP signalling in central nutrient-growth networks. Nature.

Ma, Q., Tang, R.J., Zheng, X.J., Wang, S.M., and Luan, S. (2015). The calcium sensor CBL7 modulates plant responses to low nitrate in Arabidopsis. Biochem Biophys Res Commun 468:59–65.

Marchi, L., Degola, F., Polverini, E., Terce-Laforgue, T., Dubois, F., Hirel, B., and Restivo, F.M. (2013). Glutamate dehydrogenase isoenzyme 3 (GDH3) of Arabidopsis thaliana is regulated by a combined effect of nitrogen and cytokinin. Plant Physiol Biochem 73:368–374.

Marchive, C., Roudier, F., Castaings, L., Brehaut, V., Blondet, E., Colot, V., Meyer, C., and Krapp, A. (2013). Nuclear retention of the transcription factor NLP7 orchestrates the early response to nitrate in plants. Nat Commun 4:1713.

Medici, A., and Krouk, G. (2014). The primary nitrate response: a multifaceted signalling pathway. J Exp Bot 65:5567–5576.

Medici, A., Marshall-Colon, A., Ronzier, E., Szponarski, W., Wang, R., Gojon, A., Crawford, N.M., Ruffel, S., Coruzzi, G.M., and Krouk, G. (2015). AtNIGT1/HRS1 integrates nitrate and phosphate signals at the Arabidopsis root tip. Nat Commun 6:6274.

Menz, J., Li, Z., Schulze, W.X., and Ludewig, U. (2016). Early nitrogen-deprivation responses in Arabidopsis roots reveal distinct differences on transcriptome and (phospho-) proteome levels between nitrate and ammonium nutrition. Plant J 88:717–734.

Mora-Macias, J., Ojeda-Rivera, J.O., Gutierrez-Alanis, D., Yong-Villalobos, L., Oropeza-Aburto, A., Raya-Gonzalez, J., Jimenez-Dominguez, G., Chavez-Calvillo, G., Rellan-Alvarez, R., and Herrera-Estrella, L. (2017). Malate-dependent Fe accumulation is a critical checkpoint in the root developmental response to low phosphate. Proc Natl Acad Sci U S A 114:E3563–E3572.

Muller, J., Toev, T., Heisters, M., Teller, J., Moore, K.L., Hause, G., Dinesh, D.C., Burstenbinder, K., and Abel, S. (2015). Iron-dependent callose deposition adjusts root meristem maintenance to phosphate availability. Dev Cell 33:216–230.

O'Brien, J.A., Vega, A., Bouguyon, E., Krouk, G., Gojon, A., Coruzzi, G., and Gutierrez, R.A. (2016). Nitrate Transport, Sensing, and Responses in Plants. Mol Plant 9:837–856.

O'Malley, R.C., Huang, S.S., Song, L., Lewsey, M.G., Bartlett, A., Nery, J.R., Galli, M., Gallavotti, A., and Ecker, J.R. (2016). Cistrome and Epicistrome Features Shape the Regulatory DNA Landscape. Cell 165:1280–1292.

Obertello, M., Krouk, G., Katari, M.S., Runko, S.J., and Coruzzi, G.M. (2010). Modeling the global effect of the basic-leucine zipper transcription factor 1 (bZIP1) on nitrogen and light regulation in Arabidopsis. BMC Syst Biol 4:111.

Ohkubo, Y., Tanaka, M., Tabata, R., Ogawa-Ohnishi, M., and Matsubayashi, Y. (2017). Shoot-to-root mobile polypeptides involved in systemic regulation of nitrogen acquisition. Nat Plants 3:17029.

Okamoto, M., Vidmar, J.J., and Glass, A.D. (2003). Regulation of NRT1 and NRT2 gene families of Arabidopsis thaliana: responses to nitrate provision. Plant Cell Physiol. 44:304–317.

Orozco-Cardenas, M.L., Narvaez-Vasquez, J., and Ryan, C.A. (2001). Hydrogen peroxide acts as a second messenger for the induction of defense genes in tomato plants in response to wounding, systemin, and methyl jasmonate. Plant Cell 13:179–191.

Para, A., Li, Y., Marshall-Colon, A., Varala, K., Francoeur, N.J., Moran, T.M., Edwards, M.B., Hackley, C., Bargmann, B.O., Birnbaum, K.D., et al. (2014). Hit-and-run transcriptional control by bZIP1 mediates rapid nutrient signaling in Arabidopsis. Proc Natl Acad Sci U S A 111:10371–10376.

Puga, M.I., Mateos, I., Charukesi, R., Wang, Z., Franco-Zorrilla, J.M., de Lorenzo, L., Irigoyen, M.L., Masiero, S., Bustos, R., and Rodriguez, J., et al. (2014). SPX1 is a phosphate-dependent inhibitor of PHOSPHATE STARVATION RESPONSE 1 in Arabidopsis. Proc Natl Acad Sci U S A 111:14947–14952.

Rahayu, Y.S., Walch-Liu, P., Neumann, G., Romheld, V., von Wiren, N., and Bangerth, F. (2005). Root-derived cytokinins as long-distance signals for NO_3_^-^-induced stimulation of leaf growth. J Exp Bot 56:1143–1152.

Ristova, D., Carre, C., Pervent, M., Medici, A., Kim, G.J., Scalia, D., Ruffel, S., Birnbaum, K., Lacombe, B., Busch, W., et al. (2016). Combinatorial interaction network of transcriptomic and phenotypic responses to nitrogen and hormones in the Arabidopsis thaliana root. Sci Signal 9.

Riveras, E., Alvarez, J.M., Vidal, E.A., Oses, C., Vega, A., and Gutierrez, R.A. (2015). The Calcium Ion Is a Second Messenger in the Nitrate Signaling Pathway of Arabidopsis. Plant Physiol 169:1397–1404.

Rubin, G., Tohge, T., Matsuda, F., Saito, K., and Scheible, W.R. (2009). Members of the LBD family of transcription factors repress anthocyanin synthesis and affect additional nitrogen responses in Arabidopsis. Plant Cell 21:3567–3584.

Ruffel, S., Krouk, G., Ristova, D., Shasha, D., Birnbaum, K.D., and Coruzzi, G.M. (2011). Nitrogen economics of root foraging: transitive closure of the nitrate-cytokinin relay and distinct systemic signaling for N supply vs. demand. Proc Natl Acad Sci U S A 108:18524–18529.

Ruffel, S., Poitout, A., Krouk, G., Coruzzi, G.M., and Lacombe, B. (2015). Long-distance nitrate signaling displays cytokinin dependent and independent branches. J Integr Plant Biol.

Safi, A., Medici, A., Szponarski, W., Ruffel, S., Lacombe, B., and Krouk, G. (2017). The world according to GARP transcription factors. Curr Opin Plant Biol 39:159–167.

Scheible, W.R., Gonzalez-Fontes, A., Lauerer, M., Muller-Rober, B., Caboche, M., and Stitt, M. (1997). Nitrate Acts as a Signal to Induce Organic Acid Metabolism and Repress Starch Metabolism in Tobacco. Plant Cell 9:783–798.

Shen, B., Jensen, R.G., and Bohnert, H.J. (1997a). Increased resistance to oxidative stress in transgenic plants by targeting mannitol biosynthesis to chloroplasts. Plant Physiol 113:1177–1183.

Shen, B., Jensen, R.G., and Bohnert, H.J. (1997b). Mannitol Protects against Oxidation by Hydroxyl Radicals. Plant Physiol 115:527–532.

Shin, R., Berg, R.H., and Schachtman, D.P. (2005). Reactive oxygen species and root hairs in Arabidopsis root response to nitrogen, phosphorus and potassium deficiency. Plant Cell Physiol 46:1350–1357.

Shin, R., and Schachtman, D.P. (2004). Hydrogen peroxide mediates plant root cell response to nutrient deprivation. Proc Natl Acad Sci U S A 101:8827–8832.

Stitt, M. (1999). Nitrate regulation of metabolism and growth. Curr. Opin. Plant Biol. 2:178–186.

Sutton, M.A., Oenema, O., Erisman, J.W., Leip, A., Grinsven, H.V., and Winiwarter, W. (2011). Too much of a good thing. Nature 472:159–161.

Tabata, R., Sumida, K., Yoshii, T., Ohyama, K., Shinohara, H., and Matsubayashi, Y. (2014). Perception of root-derived peptides by shoot LRR-RKs mediates systemic N-demand signaling. Science 346:343–346.

Tsukagoshi, H., Busch, W., and Benfey, P.N. (2010). Transcriptional regulation of ROS controls transition from proliferation to differentiation in the root. Cell 143:606–616.

Voegele, R.T., Hahn, M., Lohaus, G., Link, T., Heiser, I., and Mendgen, K. (2005). Possible roles for mannitol and mannitol dehydrogenase in the biotrophic plant pathogen Uromyces fabae. Plant Physiol 137:190–198.

Wang, R., Tischner, R., Gutierrez, R.A., Hoffman, M., Xing, X., Chen, M., Coruzzi, G., and Crawford, N.M. (2004). Genomic analysis of the nitrate response using a nitrate reductase-null mutant of Arabidopsis. Plant Physiol 136:2512–2522.

Wang, R., Xing, X., Wang, Y., Tran, A., and Crawford, N.M. (2009). A genetic screen for nitrate regulatory mutants captures the nitrate transporter gene NRT1.1. Plant Physiol 151:472–478.

Wu, C., Feng, J., Wang, R., Liu, H., Yang, H., Rodriguez, P.L., Qin, H., Liu, X., and Wang, D. (2012). HRS1 acts as a negative regulator of abscisic acid signaling to promote timely germination of Arabidopsis seeds. PLoS One 7:e35764.

Xu, N., Wang, R., Zhao, L., Zhang, C., Li, Z., Lei, Z., Liu, F., Guan, P., Chu, Z., Crawford, N.M., et al. (2016). The Arabidopsis NRG2 Protein Mediates Nitrate Signaling and Interacts with and Regulates Key Nitrate Regulators. Plant Cell 28:485–504.

Zhang, X., Henriques, R., Lin, S.S., Niu, Q.W., and Chua, N.H. (2006). Agrobacterium-mediated transformation of Arabidopsis thaliana using the floral dip method. Nat Protoc 1:641–646.

Zhao, M., Ding, H., Zhu, J.K., Zhang, F., and Li, W.X. (2011). Involvement of miR169 in the nitrogen-starvation responses in Arabidopsis. New Phytol 190:906–915.

